# Minute-scale coupling of chromatin marks and transcriptional bursts

**DOI:** 10.64898/2026.02.08.704500

**Authors:** Xiaohui Gao, Chaebeen Ko, Yuanchao Dong, Takeru Fujii, Satoshi Uchino, Yoshiaki Kobayashi, Akihito Harada, Hiroaki Ohishi, Yasuyuki Ohkawa, Hiroshi Kimura, Hiroshi Ochiai

## Abstract

Histone modifications are often described as stable epigenetic marks that contribute to maintaining gene-expression programs during development and environmental responses ^1–5^. However, transcription of many genes is intermittent, switching between transcriptionally active and inactive episodes within minutes ^6–10^. Whether chromatin marks around individual genes change on these rapid timescales remains unclear. Here we show that local chromatin modification signals around endogenous genes in mouse embryonic stem cells fluctuate reversibly with transcriptional state, using live imaging of individual genes together with fluorescent probes that report histone modifications^11–16^. Activation-associated acetylation and methylation marks increased in association with transcriptional activation and decreased with inactivation, whereas a Polycomb-associated repressive mark behaved oppositely. Transcriptional coactivators and both histone acetyltransferase and deacetylase complexes were enriched during transcriptionally active state, consistent with opposing enzymatic activities shaping local acetylation levels ^17,18^. Inhibiting histone deacetylases altered the durations of active and inactive events, supporting a role for deacetylation in regulating transcriptional state transitions. Thus, histone modifications undergo reversible, minute-scale changes coupled to transcriptional activity. This framework helps explain how stochastic transcriptional bursts can occur with stable gene regulation over longer timescales.

## Introduction

During development and differentiation in multicellular organisms, transcription factors and cofactors recruit chromatin-modifying enzymes to reshape local histone modification states alongside changes in gene-expression programs ^1–3^. Histone acetyltransferases (HATs), for example, deposit acetylation marks that are associated with increased chromatin accessibility and transcription, whereas histone deacetylases (HDACs) remove acetyl groups and can counteract these effects ^4,5^. Some histone modifications are relatively stable and change over hours to days, often governed by development and differentiation program ^4,5^.

Transcription at individual genes is frequently intermittent, switching between transcriptionally active and inactive states on the timescale of minutes—a behavior known as transcriptional bursting. Hereafter, we refer to these as the Active and Inactive states, respectively ^6–8^. Bursting influences both mean expression levels and cell-to-cell variability and has been observed across organisms and cell types, from single-gene measurements to genome-wide analyses ^9,10^. A growing body of work indicates that burst kinetics are shaped by dynamic enhancer–promoter interactions and by transient binding of transcription factors and coactivators that promote RNA polymerase II (RNAPII) recruitment and productive elongation ^19–23^. For example, a coactivator BRD4 binds to acetylated histones via its bromodomain and can promote transcription by recruiting the elongation factor P-TEFb ^24^, linking acetylated chromatin to transcriptional output. Indeed, single-gene imaging studies have reported preferential accumulation of BRD4 around Active loci ^11,25^. These observations raise the possibility that histone acetylation—and potentially other promoter-proximal marks—changes dynamically on the minute timescale of transcriptional bursting. However, it remains unresolved whether histone acetylation—and potentially other promoter-proximal marks—undergo rapid and reversible changes as loci enter and exit the Active state, or whether histone modification states are comparatively stable and only indirectly related to bursting dynamics.

To address this question, we combined STREAMING-tag live-cell single-gene imaging with fluorescent probes that report histone modifications and RNAPII phosphorylation^11–16^. Using multiple endogenous genes in mouse embryonic stem cells (mESCs), we quantified locus-centered chromatin signals across the Active/Inactive cycle. We further focused on histone acetylation and tested how opposing enzymatic activities—including the HAT p300 and HDACs—relate to transcriptional state transitions.

## Results

### Histone-mark signals at endogenous loci fluctuate with transcriptional state during bursting

To test whether chromatin marks change on the timescale of transcriptional bursting, we analyzed histone modifications at endogenous *Nanog*, *Sox2*, *Usp5*, and *Dnmt3L* loci in mouse embryonic stem cells (mESCs) using STREAMING-tag system^11^ (Fig. 1a and Supplementary Fig. 1). The STREAMING-tag contains MS2 and TetO repeats, enabling simultaneous visualization of nascent transcripts by an MS2 coat protein fused to cyan fluorescent protein (MCP-CFP) and the tagged locus by a mutant Tet repressor fused to yellow fluorescent protein (mTetR-YFP) in living cells. Based on the presence or absence of a locus-proximal MCP focus, we classified the gene state at each time point as Active or Inactive states, which allows detecting the state switching at individual loci in time (Fig. 1a, Methods)^11^.

**Figure 1.**
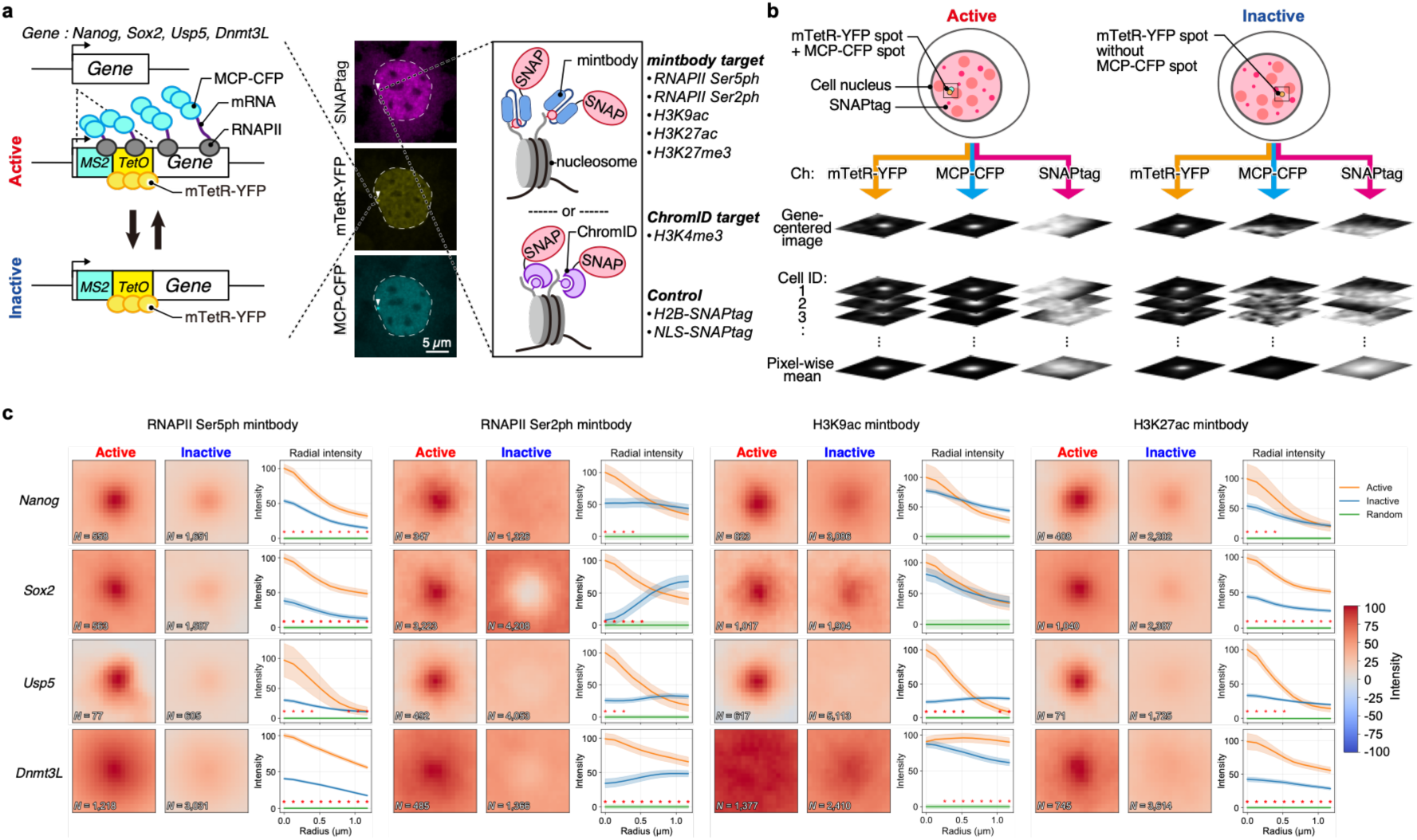
Locus-centered signals of histone modification and phospho-RNAPII around genes change during the transcriptional-burst cycle. **(a)** Schematic of the STREAMING-tag single-gene imaging system. In target genes (*Nanog*, *Sox2*, *Usp5*, *Dnmt3L*), the STREAMING-tag cassette is knocked in, and moderate expression of MCP-CFP and mTetR-YFP enables visualization of the gene locus and transcriptional activity. Co-expression of mintbody or ChromID probes fused to a SNAPtag allows simultaneous visualization of histone modifications. Arrowheads mark the locus; the MCP-CFP focus co-localizes with the mTetR-YFP locus focus during transcriptionally Active frames. **(b)** Schematic of locus-centered quantification of histone-mark and phospho-RNAPII signals at endogenous loci. **(c)** Quantitative analysis of locus-centered signals reported by histone mark and phospho-RNAPII probes. For each locus instance, a 19 × 19-pixel crop (0.13 µm per pixel) centered on the mTetR focus (z-plane of the focus) was extracted from each channel and grouped by transcriptional state (Active versus Inactive states). Pixel-wise mean locus-centered images were computed for each state after subtracting the mean image of size-matched random nuclear control crops sampled away from the locus. Radial intensity profiles show the mean ± s.e.m. (standard error of the mean) across locus instances of the background-subtracted intensity in concentric 1-pixel annuli around the center. Red asterisks indicate radii with significant differences between Active and Inactive states (two-sided Mann-Whitney U test at each radius with Benjamini-Hochberg correction across radii; *q* < 0.05).

To visualize the RNAPII and chromatin mark states in the same cells, we expressed SNAPtag fusions of modification-specific probes based on either a modification-specific intracellular antibody (mintbody) or a binding module for chromatin-dependent protein identification (ChromID). These probes reported RNAPII C-terminal domain (CTD) phosphorylation (Ser5ph and Ser2ph) and histone H3 modifications associated with active chromatin—H3K9ac (H3 Lys9 acetylation), H3K27ac (H3 Lys27 acetylation) and H3K4me3 (H3 Lys4 trimethylation)—or Polycomb repression (H3K27me3; H3 Lys27 trimethylation)^12–15^. Because these probes bind reversibly to the target modifications with residence times on the order of a few to tens of seconds, minute-scale kinetics can be analyzed, and their expression has minimal effects on cell cycle progression, differentiation, and animal development^26,27^. SNAPtag fusions with a nuclear localization signal (NLS-SNAPtag) or histone H2B (H2B-SNAPtag) served as reference reporters for nuclear distribution and local chromatin density/compaction, respectively (Supplementary Fig. 2). To minimize variability arising from transgene dosage, we isolated populations with moderate and relatively uniform mTetR, MCP, and SNAPtag expression by fluorescence-activated cell sorting (FACS). Using these cells, we quantified locus-centered signals by cropping mTetR-centered areas and then drawing radial profiles in Active and Inactive frames (Fig. 1b; Supplementary Fig. 2–3; Methods).

Active frames showed strong enrichment of RNAPII Ser5ph and Ser2ph at all tagged loci analyzed (Fig. 1c). Similarly, when quantified in the locus-centered core pixel region, three active histone-marks, i.e., H3K9ac, H3K27ac and H3K4me3, were significantly higher in Active frames than during Inactive frames except H3K9ac at *Nanog*, *Sox2*, and *Dnmt3L* loci, whereas H3K27me3 showed the opposite tendency, with relatively higher signals in Inactive frames (Fig. 1c and Supplementary Fig. 4a). H2B-SNAPtag tended to be more concentrated at the loci in Inactive than in Active states, whereas NLS-SNAPtag showed little difference in the two states (Supplementary Fig. 4a). Notably, H3K27ac and H3K27me3—mutually exclusive modifications on H3K27—displayed reciprocal state dependence, which argues against a uniform change in probe access to more open chromatin in Active state and instead supporting mark-specific differences in local epitope availability during state switching.

To assess cell-to-cell variability, we quantified per-locus probe intensities at the tagged loci (Supplementary Fig. 4b). Probe intensities were highly variable across cells in both states and thus could not reliably predict transcriptional state, although weak positive correlations for RNAPII Ser5ph/Ser2ph and H3K27ac and weak negative correlations for H3K27me3 and H2B were observed (Supplementary Fig. 4c). In addition, reanalysis of ChIP-seq tracks confirmed that (except for H3K27me3) these features are enriched around the loci analyzed here relative to the genome-wide average (Supplementary Fig. 5). Together, these results indicate that locus-centered signals for chromatin marks fluctuate with transcriptional state switching on minute timescales.

We next asked whether state-dependent enrichment extends beyond chromatin marks to transcription-related regulators in an orthogonal fixed-cell dataset (Fig. 2a)^25^.

**Fig. 2.**
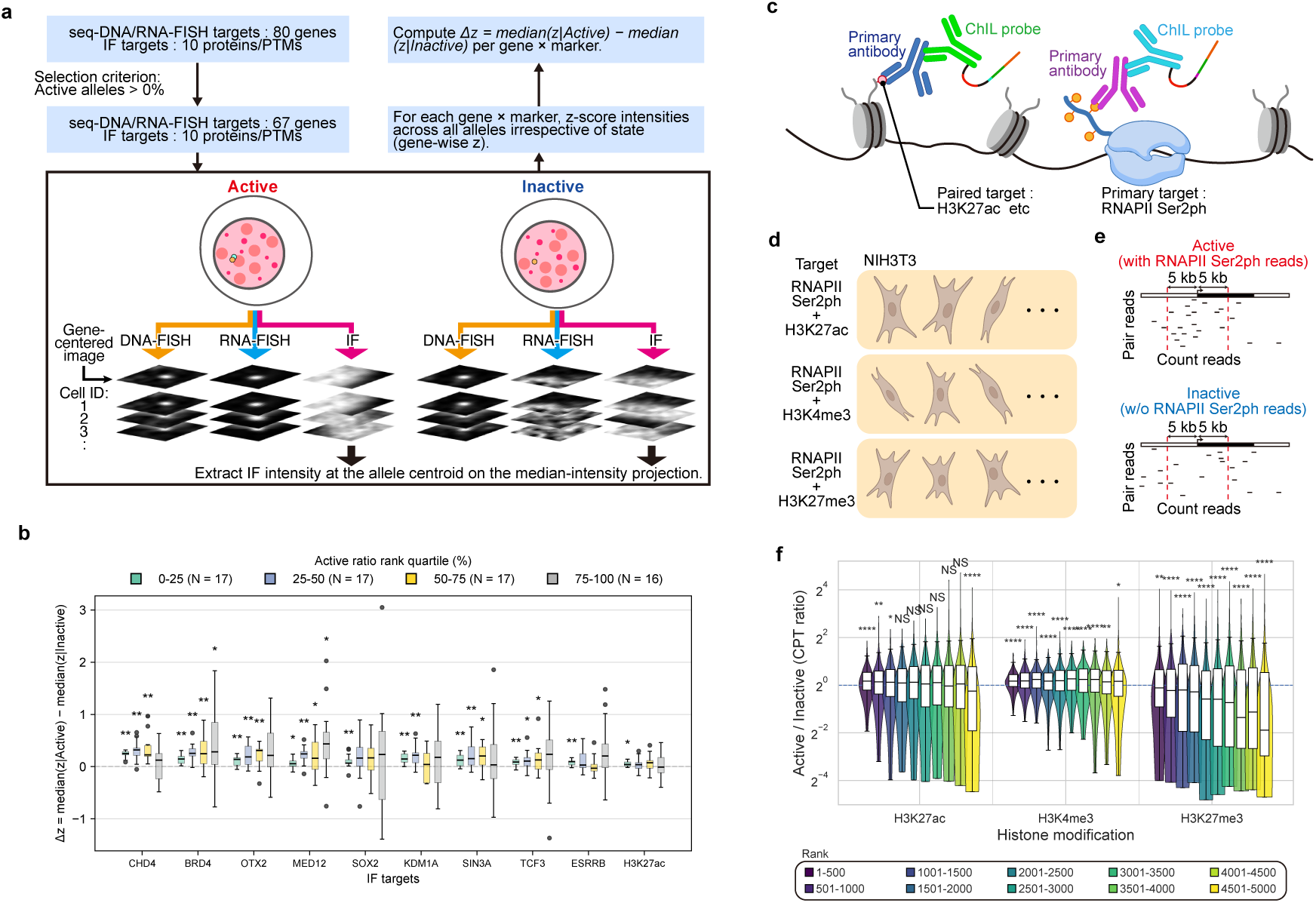
Enrichment trends of histone modifications and transcription-related factors around genes during the transcription-burst cycle in multimodal seq-DNA/RNA/IF-FISH data and sci-mtChIL-seq data. **(a)** Workflow of reanalysis of multimodal seq-DNA/RNA/IF-FISH data in mouse embryonic stem cells (mESCs)^25^. For each allele identified by seq-DNA-FISH, transcriptional state was classified as Active or Inactive states using the corresponding seq-RNA-FISH signal. Immunostaining intensity for each target (10 proteins/PTMs) was extracted at the allele centroid from the matched 3D IF stack and standardized as a gene-wise z score. For each gene and target, Δ*z* was computed as *median*(*z*|*Active*) − *median*(*z*|*Inactive*). Genes with at least one Active allele (Active alleles >0%) and sufficient Active/Inactive instances were retained (67 genes) and ranked by their Active-allele ratio; genes were then stratified into activity-rank quartiles. **(b)** Distributions of Δ*z* values for each immunostaining marker across genes, shown separately for the activity-rank quartiles (0–25%, 25–50%, 50–75% and 75–100%; *n* = 17, 17, 17 and 16 genes, respectively). Box plots indicate median and interquartile range (IQR); whiskers denote 1.5× IQR. Points represent individual genes. Asterisks indicate Benjamini–Hochberg FDR-adjusted *q* values for two-sided one-sample Wilcoxon signed-rank tests against 0 within each marker × quartile. (* *q* ≤ 0.05, ** *q* ≤ 0.01). **(c)** Schematic of sci-mtChIL-seq, in which RNAPII Ser2ph is used as a primary target and histone modifications are measured as paired targets within the same single cells. **(d)** Overview of the published NIH3T3 sci-mtChIL-seq datasets analyzed here, in which RNAPII Ser2ph was paired with H3K27ac, H3K4me3, or H3K27me3^28^. **(e)** Definition of read counts for targets paired with RNAPII Ser2ph. A gene was defined as Active state if RNAPII Ser2ph reads were detected within 1 kb upstream and 5 kb downstream of the TSS. Reads for the paired target were counted within TSS ±5 kb, normalized as counts per thousand (CPT), and compared between Active and Inactive states. Analyses were restricted to an RNAPII-covered set (top 5,000 genes by RNAPII Ser2ph counts). **(f)** Distributions of gene-wise log_2_(Active/Inactive) CPT ratios for the paired histone modifications across activity-frequency bins. Genes in the RNAPII-covered set were ranked by a ChIL-based activity frequency score (fraction of cells classified as Active state) and partitioned into consecutive 500-gene bins (ranks 1–500 to 4501–5000; colors). Violin plots show distributions and embedded box plots indicate medians and interquartile ranges; the dashed horizontal line indicates a ratio of 1 (log_2_ ratio = 0). Statistical significance of deviation of log_2_(Active/Inactive) from 0 was assessed for each (target, bin) using a two-sided one-sample Wilcoxon signed-rank test, followed by Benjamini–Hochberg correction across all comparisons; asterisks denote adjusted significance (NS, *q* ≥ 0.05; * *q* < 0.05; ** *q* < 0.01; *** *q* < 0.001; **** *q* < 1×10^-4^).

Reanalysis of published multimodal seq-DNA/RNA/IF-FISH data in mESCs showed that alleles classified as transcriptionally Active state tend to exhibit higher local immunofluorescence signals for transcription-related factors as well as H3K27ac at the corresponding gene loci (Fig. 2b; Supplementary Fig. 6a). Particularly for BRD4 and MED12, pluripotency-related transcription factors (SOX2, TCF3, ESRRB and OTX2), and HDAC-associated factors (KDM1A, SIN3A and CHD4), Active-biased enrichment was most apparent among genes with the highest Active-allele fractions and progressively attenuated toward lower-activity quartiles (Fig. 2b).

Because the analyses above are limited to spatial information, we could not exclude the possibility that the apparent state-dependent signals reflect locus repositioning toward (or away from) nuclear regions enriched for specific histone marks, rather than bona fide changes in chromatin features near the gene neighborhood (for example, promoters or nearby enhancers). To complement the locus-centered imaging readouts with an orthogonal locus-resolved approach, we reanalyzed published single-cell combinatorial indexing multi-target chromatin integration labeling followed by sequencing (sci-mtChIL-seq) data in NIH3T3 cells, which profiles elongating RNAPII (Ser2ph) and a paired histone modification at the same loci in single cells, thereby providing a complementary test that is less constrained by optical resolution (Fig. 2c–e) ^28^. Using RNAPII Ser2ph occupancy to stratify gene instances into transcriptionally engaged versus unengaged groups (see Methods), we compared promoter-proximal paired-target signals across activity-frequency bins (500-gene bins) (Fig. 2e). H3K4me3 was consistently higher and H3K27me3 consistently lower in the RNAPII-positive group across bins, whereas H3K27ac was found enriched in the highest RNAPII-positive frequency bins and less-enriched for lower-frequency bins (Fig. 2f; Supplementary Fig. 6). Together, these analyses support the interpretation that transcriptional engagement is accompanied by mark-specific differences in the local chromatin environment, while indicating that the strength of coupling depends on both the histone mark and transcription activity.

### Temporal coupling of acetylation regulators to transcriptional state switching

The transcription state-dependent fluctuations of acetylation-associated marks suggest that acetylation around loci is regulated on minute timescales. We therefore asked whether proteins that write, read, or erase acetylation—and their associated complexes—co-accumulate with transcriptional state switching during bursting.

Genome-wide chromatin-state analysis in mESCs indicated that, alongside H3K27ac and p300, multiple HDACs are enriched at active promoters and enhancers (Fig. 3a–c). Because HDAC catalytic activity is typically exerted within multiprotein assemblies, we focused on HDAC1/2-associated modules (including NuRD and SIN3 complexes) and on the HDAC3–NCoR/SMRT module, and therefore included representative complex components (CHD4, SIN3A and NCOR2 (also known as SMRT)) in subsequent analyses ^29–33^. In the seq-DNA/RNA/IF-FISH reanalysis, CHD4 and SIN3A—HDAC-interacting factors—showed Active-biased enrichment most clearly among the higher-activity genes, together with H3K27ac, BRD4, and the transcription factor SOX2 (Fig. 2b)^25^. In addition, reanalysis of the Modification Atlas of Regulation by Chromatin States (MARCS) proteomic interactome maps indicated that HDAC1/2 and HDAC-containing assemblies such as NuRD are preferentially associated with acetylated histone features (for example, H3ac) (Supplementary Fig. 7a–c)^34^. The SIN3 complex, which contains HDAC1/2, showed preferential association with H3K4me3 (Supplementary Fig. 7d), consistent with recruitment of deacetylase activity to transcription-associated chromatin contexts.

**Fig. 3.**
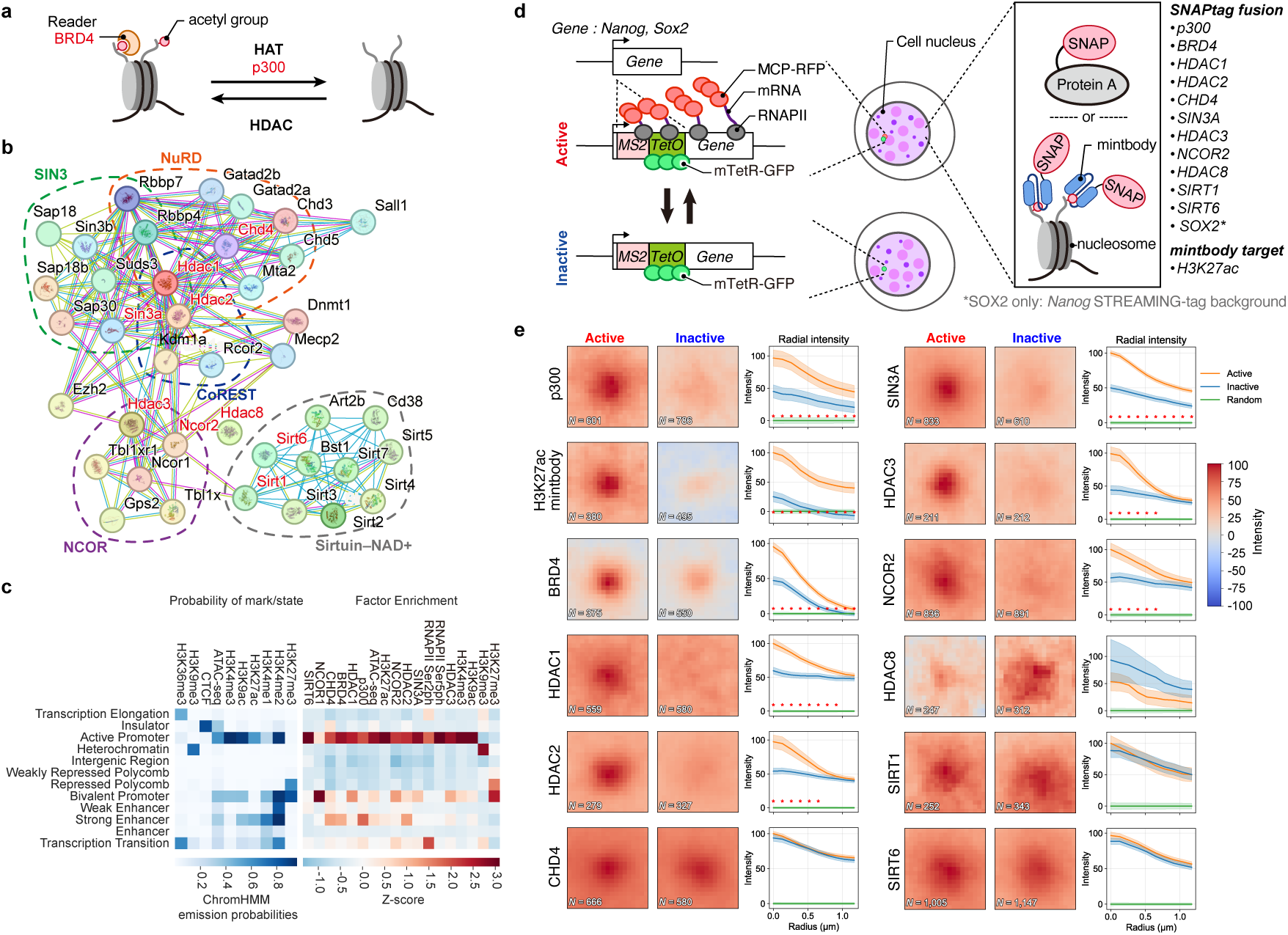
Accumulation dynamics of factors mediating the installation and removal of histone acetylation around genes. **(a)** Schematic of acetylation deposition and removal. **(b)** Interaction network of HDAC-associated factors. Protein-protein interaction networks centered on the HDAC family were visualized using the STRING database. Nodes represent proteins, and edges represent known or predicted functional associations. Edge colors indicate the types of interaction evidence (e.g., experimental evidence, curated databases, coexpression, and text mining), and densely connected clusters within the network reflect functional modules involved in transcriptional repression complexes and chromatin modification. **(c)** Chromatin state-dependent localization tendencies of histone modifications and transcription-related factors in mESCs. Twelve chromatin states were defined using ChromHMM (mm10, 12-state model based on previously published data)^43^. Left, ChromHMM emission probabilities (i.e., the probability of observing each histone mark/factor given a chromatin state). Right, relative enrichment of factors across chromatin states, quantified as Z-scores from ChIP-seq signals for the HDAC family, p300, and transcription-related factors (e.g., RNAPII Ser2ph). **(d)** Schematic of the STREAMING-tag single-gene imaging system. In target genes (*Nanog* and *Sox2*), the STREAMING-tag cassette is knocked in, and moderate expression of MCP-RFP and mTetR-GFP enables visualization of the gene locus and transcriptional activity. Co-expression of mintbody or protein of interest fused to a SNAPtag allows simultaneous visualization of gene locus, transcriptional activity, and protein of interest. SOX2-SNAPtag data were obtained only in the *Nanog*-STREAMING-tag background. **(e)** Quantitative analysis of accumulation of the indicated protein of interest or H3K27ac centered on the *Nanog* gene locus in the Active versus Inactive state. Mean locus-centered images and radial intensity profiles were computed from 19 × 19-pixel crops (0.13 µm per pixel) centered on the mTetR focus after subtracting the mean random nuclear control image. Radial profiles show mean ± s.e.m. across locus instances. Red asterisks indicate radii with significant differences between states (two-sided Mann-Whitney U test with Benjamini-Hochberg correction across radii; *q* < 0.05).

To directly measure factor behavior during bursting at single loci, we performed live-cell imaging in *Nanog*- or *Sox2*-STREAMING-tag cells expressing mTetR fused to green fluorescent protein (mTetR-GFP) and MCP fused to red fluorescent protein (MCP-RFP). We monitored SNAP-tagged factors, including endogenously tagged BRD4, SIN3A and CHD4, and exogenously expressed SNAPtag fusions of p300, HDAC1/2/3/8, SIRT1, SIRT6, and NCOR2 (Supplementary Table 1). A SNAP-tagged H3K27ac mintbody was used to monitor an acetylation readout. Endogenous SOX2 was monitored using a *Sox2*-SNAPtag knock-in line generated only in the *Nanog*-STREAMING-tag background (Fig. 3d; Supplementary Fig. 8). Imaging populations were enriched by FACS for moderate reporter expression, and nuclear mean reporter intensities showed only limited state dependence (Supplementary Fig. 9).

Snapshot analyses revealed robust enrichment of p300 and BRD4 around the locus during the Active state, consistent with the Active-state enrichment of the H3K27ac mintbody both at the *Nanog* locus (Fig. 3e) and the *Sox2* locus (Supplementary Fig. 10). Notably, HDAC1/2 and SIN3A, as well as HDAC3 and NCOR2, also tended to accumulate preferentially during Active frames (Fig. 3e and Supplementary Fig. 10). In contrast, CHD4, HDAC8, SIRT1 and SIRT6 showed no clear state dependence under the conditions tested (Fig. 3e and Supplementary Fig. 10). These results indicate that Active loci are accompanied not only by acetylation-associated writers and readers but also by selected deacetylase modules, consistent with local balancing of opposing enzymatic activities during activation. Consistent with the fixed-cell reanalysis (Fig. 2b), endogenous SOX2 also showed enrichment at the *Nanog* locus during Active frames (Fig. 4a), motivating time-resolved analyses of TF and cofactor dynamics.

**Fig. 4.**
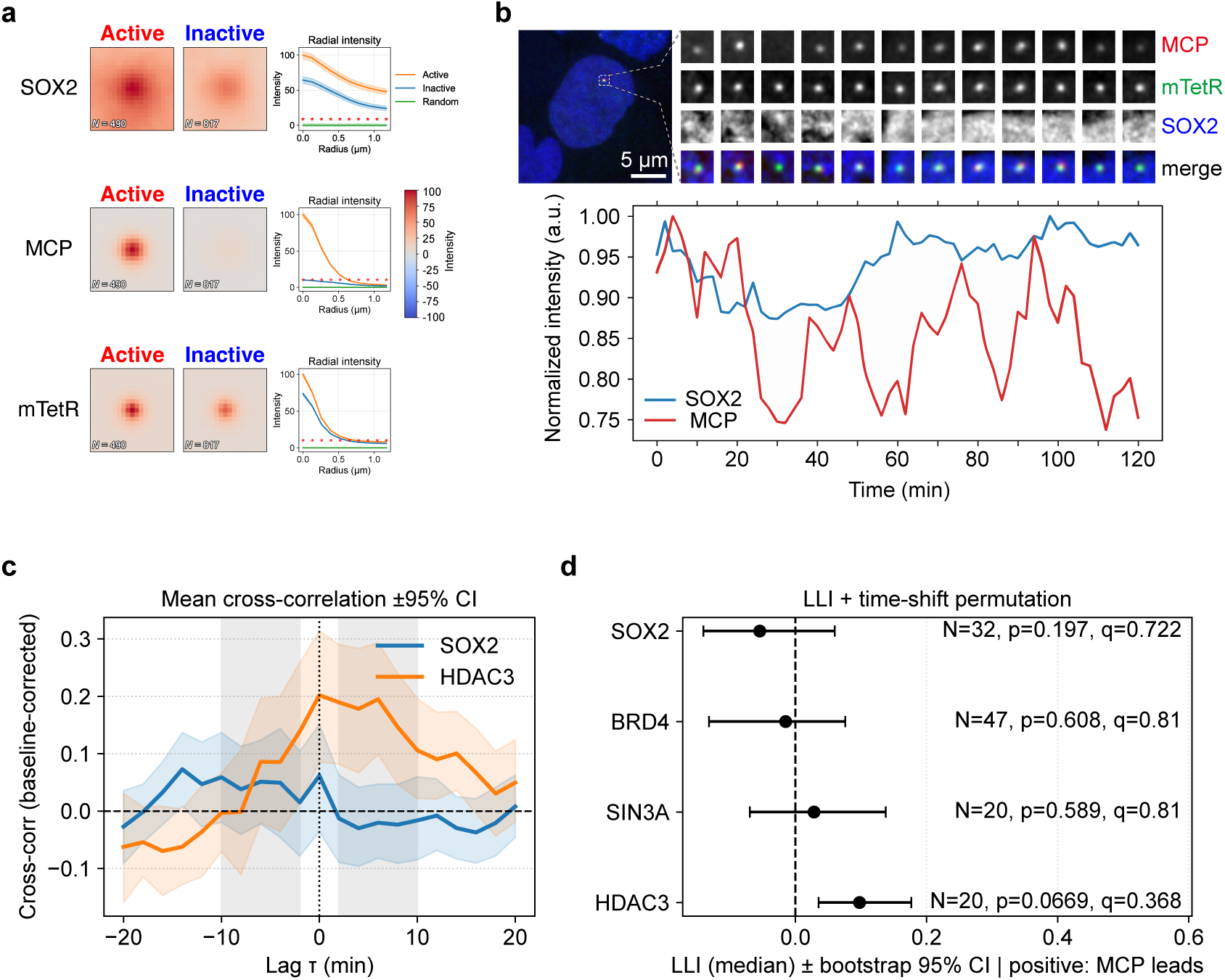
Time-resolved relationship between factor accumulation and transcriptional activity at the *Nanog* locus. **(a)** Accumulation of SOX2-SNAPtag around the *Nanog* STREAMING-tag-marked locus in *Nanog* STREAMING-tag knock-in / Sox2-SNAPtag knock-in cells expressing moderate levels of MCP-RFP and mTetR-GFP. Pixel-wise mean locus-centered images (computed from mTetR-centered crops across cells) are shown for SOX2-SNAPtag, MCP-RFP, and mTetR-GFP in transcriptionally active versus inactive frames, together with radial intensity profiles (Active, Inactive, and Random nuclear positions). **(b)** Representative time-lapse imaging of a single *Nanog* locus acquired at 2-min intervals for 2 h. Top, maximum intensity projections of MCP-RFP, mTetR-GFP, SOX2-SNAPtag, and merged images. Bottom, normalized fluorescence intensities of MCP and SOX2 quantified within the mTetR-defined region. Scale bar, 5 μm. **(c)** Mean cross-correlation functions between MCP-RFP intensity (transcriptional activity) and SOX2-SNAPtag or HDAC3-SNAPtag intensity at the locus. Trajectories were locally detrended and z-scored per cell prior to cross-correlation; cross-correlation curves were baseline-corrected by subtracting the weighted mean correlation at |lag| = 7–10 frames. Negative lag (*τ*) indicates SNAPtag leads MCP-RFP. Lines indicate the mean across cells and shaded areas indicate bootstrap 95% confidence intervals. **(d)** Lead-lag index (LLI) summarizing asymmetry of the baseline-corrected cross-correlation within ±2-10 min (excluding *τ* = 0). Points indicate the median LLI across cells and error bars indicate bootstrap 95% confidence intervals. Positive LLI indicates MCP leads. *P* values were obtained by a two-sided time-shift permutation test (within-cell circular shift), and *q* values represent Benjamini-Hochberg FDR correction across the tested factors. *N* denotes the number of cells analyzed.

Finally, to assess the temporal relationship between factor accumulation and transcriptional fluctuations, we performed time-lapse imaging at 2-min intervals for 2 h and quantified locus-centered SNAPtag and MCP signals (Fig. 4b). Cross-correlation analysis showed that SOX2, BRD4, SIN3A and HDAC3 peak near zero lag relative to transcriptional signal (Fig. 4c and Supplementary Fig. 11a). To test whether factor accumulation systematically precedes or follows transcription (i.e., whether SNAPtag signals lead or lag MCP fluctuations), we quantified asymmetry in the cross-correlation using a lead–lag index and assessed significance by within-cell time-shift permutation. This analysis with within-cell time-shift permutation tests did not support a statistically significant systematic lead or lag at this temporal resolution (Fig. 4d and Supplementary Fig. 11b), suggesting that factor accumulation is tightly coupled to transcription and is largely contemporaneous on minute timescales.

### Effect of HDAC inhibition on transcription-burst dynamics

Given the co-enrichment of HDAC modules at Active loci, we asked whether deacetylase activity contributes to transcriptional state transitions on minute timescales. We acutely inhibited HDAC activity and quantified Active and Inactive dwell times from live transcription traces. Cells were treated with vehicle (dimethyl sulfoxide; DMSO), the pan-HDAC inhibitor trichostatin A (TSA), or the HDAC3-selective inhibitor RGFP966, and *Nanog* and *Sox2* transcription was imaged at 1-min intervals for 1 h (Fig. 5a). Using the MCP signal at the mTetR-marked locus, we classified each time point into Active or Inactive states and extracted contiguous dwell times for each state (Fig. 5a,b).

**Fig. 5.**
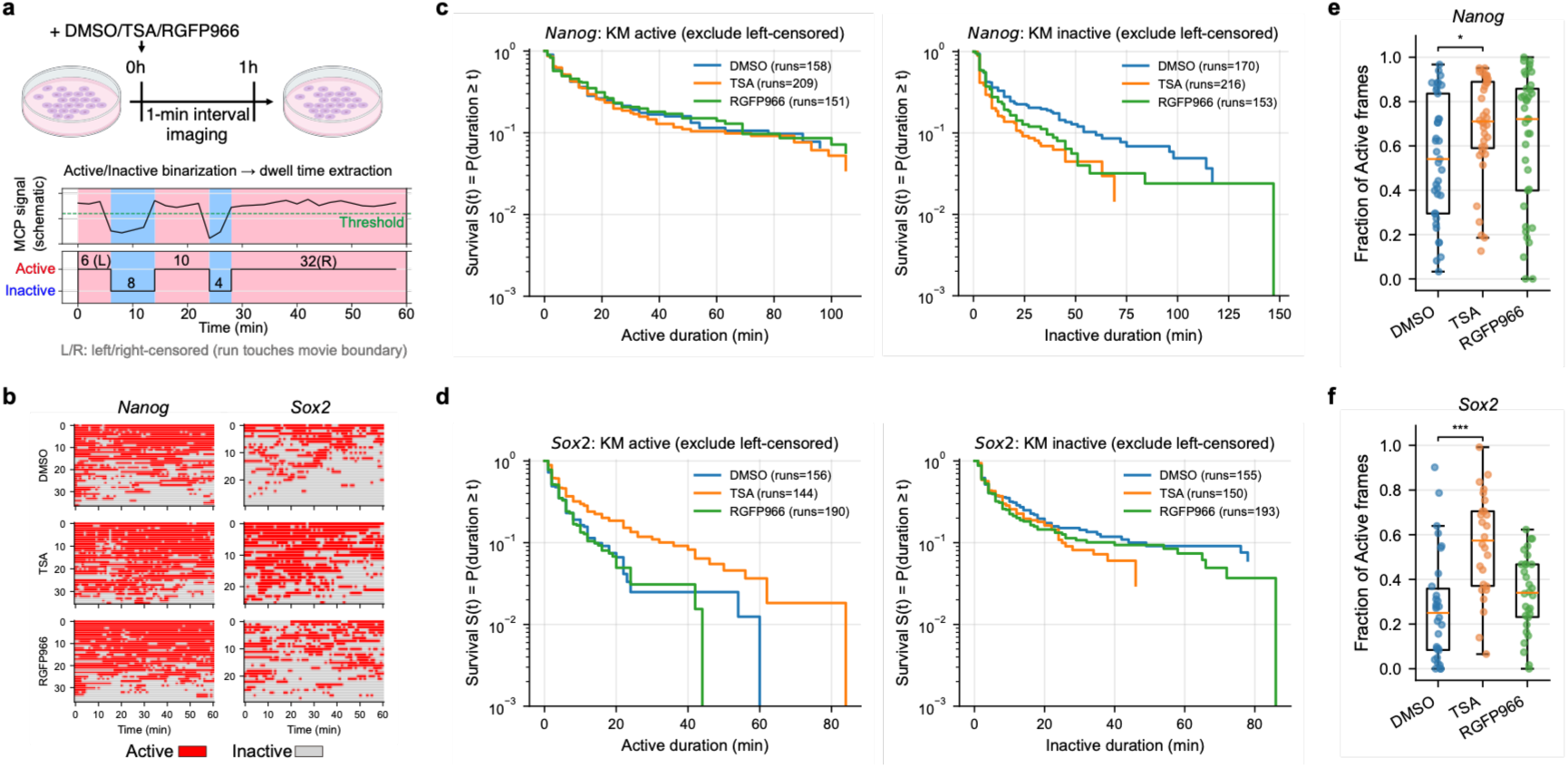
HDAC inhibition reshapes transcriptional state dynamics in a locus-dependent manner. **(a)** Experimental design and analysis schematic. Cells were treated with vehicle (DMSO), trichostatin A (TSA), or the HDAC3-selective inhibitor RGFP966 and imaged at 1-min intervals for 1 h. MCP signal at the mTetR-marked locus was converted into a binary transcriptional state (Active/Inactive) based on spot detection and temporal continuity, and contiguous runs were used to compute dwell times. Runs touching the start or end of the movie were treated as left- or right-censored, respectively (schematic). **(b)** Representative single-cell state traces (rows, individual cells; columns, time) for *Nanog* and *Sox2* under each condition. Red indicates Active frames and grey indicates Inactive frames. **(c and d)** Kaplan-Meier survival curves of Active and Inactive dwell times for *Nanog* (c) and *Sox2* (d), excluding left-censored runs. Right-censored runs were included as censored observations. Curves show the probability that a dwell time persists for at least t minutes. The number of runs analyzed per condition is indicated in each panel. **(e and f)** Cell-level duty cycle (fraction of frames classified as Active state over the 1-h movie) for *Nanog* (e) and *Sox2* (f). Points denote individual cells and box plots summarize distributions (center line, median; box, interquartile range; whiskers, 1.5× interquartile range). Statistical comparisons are two-sided Mann-Whitney U tests with Holm correction versus DMSO; asterisks denote Holm-adjusted significance (ns, *p*≥0.05; \**p* < 0.05; \*\*\**p* < 0.001).

HDAC inhibition reshaped dwell-time distributions in a locus-dependent manner (Fig. 5c–f). At the *Nanog* locus, Active-state survival curves were only modestly altered across conditions, whereas Inactive-state dwell times were shortened under TSA and showed a similar, but weaker, shift under RGFP966 (Fig. 5c). Consistent with these relatively subtle changes, the duty cycle (fraction of frames classified as Active state) shifted modestly in TSA, and not significantly in RGFP966, compared with DMSO (Fig. 5e).

At the *Sox2* locus, TSA increased the prevalence of prolonged Active episodes and concurrently reduced Inactive-state dwell times (Fig. 5d), resulting in a marked increase in duty cycle (Fig. 5f). RGFP966 produced more modest effects: it did not significantly change duty cycle, but its influence was more evident in the distribution of Inactive-state dwell times than in Active-state persistence (Fig. 5d,f). Together, these results indicate that HDAC activity can tune transcriptional state transitions during bursting on minute timescales, with distinct and locus-dependent responses to pan-HDAC versus HDAC3-selective inhibition.

## Discussion

Chromatin-based gene regulation is often discussed in terms of relatively stable epigenetic states ^1–5^, whereas transcription at many genes is intermittent and switches on minute timescales ^6–10^. Our live-cell measurements bridge this gap by showing that locus-centered chromatin-mark–associated signals around endogenous genes can fluctuate reversibly within minutes in step with transcriptional state switching. Activating features, including histone acetylation and H3K4me3, tend to rise during Active episodes and decline during Inactive episodes, whereas a Polycomb-associated repressive mark shows the opposite tendency. Together with the co-enrichment of acetylation writers, readers and selected deacetylase modules at Active loci, these findings support a view in which bursts unfold within a dynamically balanced chromatin environment, providing a mechanistic route by which rapid transcriptional fluctuations can be coupled to longer-term regulatory stability (Fig. 6).

**Fig. 6.**
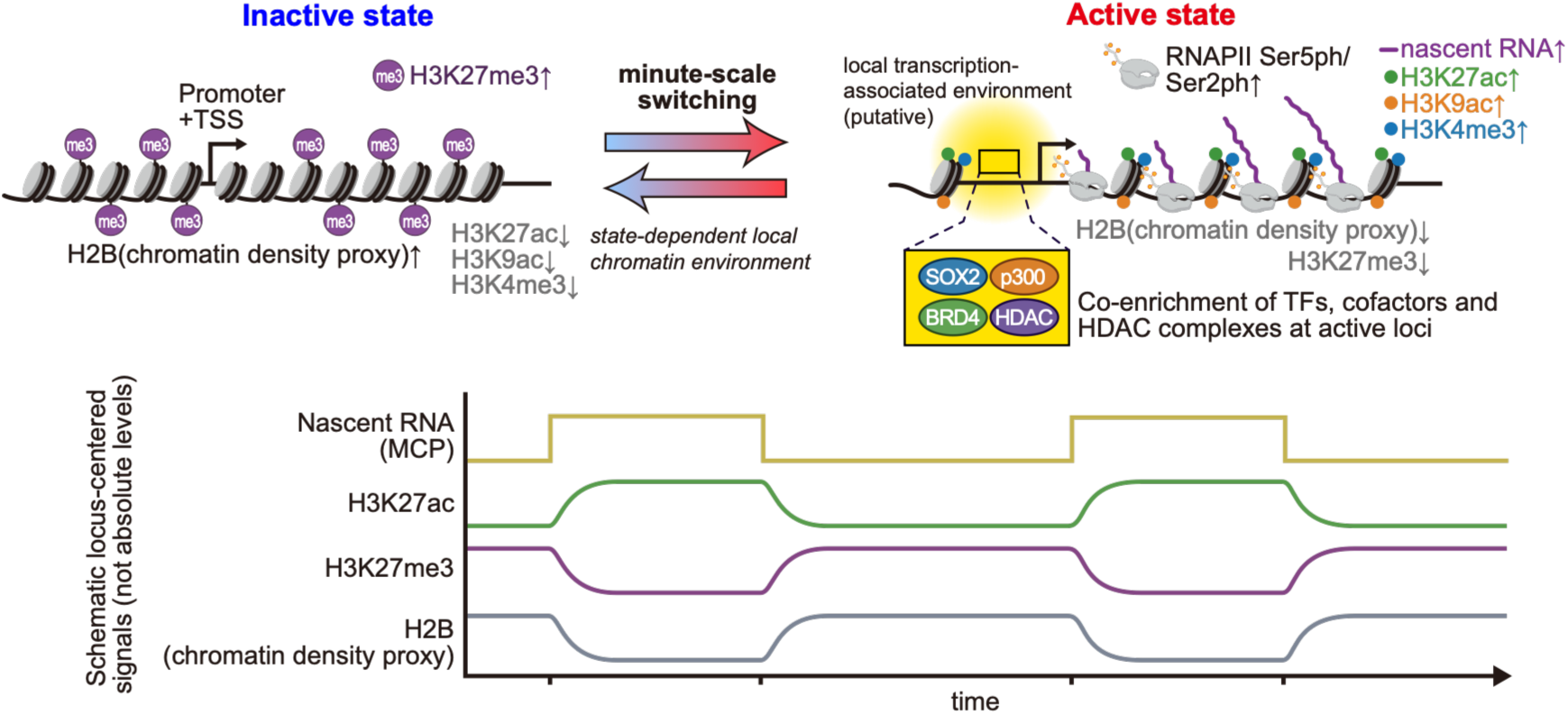
Working model for minute-scale coupling between transcriptional state switching and locus-centered chromatin signals. Schematic summary of the live-cell imaging results in mouse embryonic stem cells. Endogenous gene loci were monitored at single loci and classified into Active and Inactive states based on the presence or absence of nascent RNA signal (MCP) at the locus. During the Active state, locus-centered signals for RNAPII CTD phosphorylation (Ser5ph/Ser2ph) and activating chromatin marks (H3K27ac, H3K9ac and H3K4me3) are elevated, and transcription-associated factors/cofactors and HDAC complexes co-enrich at Active loci (illustrated by SOX2, p300, BRD4 and HDAC complexes). During the Inactive state, locus-centered signals for H3K27me3 and H2B (chromatin density proxy) are relatively higher, could reflect a more compact local chromatin environment. The bidirectional arrow indicates minute-scale switching between Active and Inactive states associated with state-dependent local chromatin environments. Bottom, schematic time traces illustrate representative locus-centered signals across switching cycles. Signals represent locus-centered probe enrichment/epitope availability and are not absolute modification levels; traces are schematic and not to scale and do not imply a specific causal order.

A key interpretive concern for locus-centered fluorescence measurements is whether state-dependent signals could arise from locus repositioning toward nuclear regions enriched for specific marks rather than from changes in the local chromatin neighborhood. Although diffraction-limited imaging cannot fully exclude repositioning effects, two orthogonal fixed-cell reanalyses support an allele- and locus-resolved association between transcriptional state and chromatin features. In multimodal seq-DNA/RNA/IF-FISH data, multiple transcription-related factors and chromatin features show Active-biased enrichment most clearly among higher-activity genes, with attenuation toward lower-activity bins as Active instances become sparse^25^. Likewise, sci-mtChIL-seq analyses reveal mark-specific differences between Active and Inactive gene states, including higher H3K4me3 and lower H3K27me3 at Active genes, while the strength of the H3K27ac shift depends on gene activity and gene subset. These observations are consistent with genuine locus-to-locus variability in coupling, and they also highlight technical constraints inherent to sparse single-cell epigenomic measurements, including false-negative Active assignments when RNAPII counts are low and ambiguity in allele attribution in a near-tetraploid/aneuploid background.

What molecular mechanisms could generate minute-scale, reversible mark changes during bursting? Histone acetylation is installed by HATs and removed by HDACs, and increasing evidence suggests that active regulatory elements can be governed by a dynamic balance of opposing enzymatic activities rather than by unidirectional “writing” followed by “erasing” ^17,18^. Consistent with this framework, our chromatin-state analyses indicate that multiple HDACs occupy active promoters and enhancers alongside p300 and H3K27ac. Proteomic interaction maps further support the capacity of HDAC-containing assemblies (including NuRD- and SIN3-related modules and HDAC3–NCOR2/SMRT complexes) to associate with acetylated and transcription-associated chromatin contexts ^34^. At the single-locus level, the co-enrichment of p300 and BRD4 together with selected HDAC modules during Active episodes suggests a locally coupled circuit in which activation-associated activities promote acetylation and engagement of the transcription machinery, while co-recruited deacetylase complexes restrain acetylation and thereby tune the duration of Active episodes and transitions back to the Inactive state.

A related question is whether chromatin-mark changes and factor recruitment precede transcriptional activation or instead track it contemporaneously. Live-cell studies using histone-modification probes in engineered reporter-array systems have suggested that H3K27ac can rise ahead of transcriptional activation, consistent with chromatin priming in some contexts^35,36^. However, array-based reporters integrate signals across tandem repeats and differ in locus architecture and temporal readouts from endogenous single-locus bursting, and therefore gene structure and sampling rate may influence the apparent ordering. In our endogenous-locus measurements at two-minute resolution, SOX2, BRD4, SIN3A and HDAC3 signals were tightly coupled to transcriptional signal, with cross-correlation peaks near zero lag and no statistically supported systematic lead–lag asymmetry. Any temporal ordering may therefore be subtle or occur on sub-minute timescales, potentially reflecting rapid multi-step recruitment events that are blurred by minute-level sampling. Higher temporal resolution, additional endogenous tagging and perturbations that selectively disrupt specific interaction interfaces will be necessary to establish causality and ordering.

In this context, SOX2 is of particular interest because it has been reported to associate with HDAC-containing complexes and to link to p300-containing coactivator activity ^37,38^, potentially enabling coordinated engagement of activating and restraining enzymatic modules. Likewise, BRD4 contributes to productive elongation through regulation of P-TEFb and pause release ^24,39^, and proteomic and functional studies suggest that BET proteins can interface with chromatin-remodeling and corepressor complexes in other contexts ^40–42^. Together, these connections suggest testable routes by which a TF/cofactor hub could engage both activating and restraining modules during bursts.

Functional perturbations support a role for deacetylation in shaping transcriptional state transitions. Acute HDAC inhibition reshaped the dwell-time distributions of Active and Inactive states in a locus-dependent manner, consistent with deacetylation contributing to the establishment and/or maintenance of the Inactive state and tuning burst kinetics on minute timescales. The distinct responses of *Nanog* and *Sox2* to broad versus HDAC3-selective inhibition further point to locus-specific regulatory architectures and chromatin contexts as determinants of how the writer–reader–eraser balance is implemented at individual genes.

Several limitations warrant consideration. First, fluorescent probes report local concentration and accessibility of modification epitopes within the optical point-spread function rather than absolute modification stoichiometry, and microscopy cannot fully separate local neighborhood changes from modest repositioning. Second, binary state calling captures switching but discards amplitude information and may miss brief events. Third, locus coverage and factor coverage remain limited. Extending endogenous tagging, increasing temporal resolution, and integrating targeted perturbations of writer/reader/eraser modules should help delineate the molecular feedbacks that link chromatin dynamics to burst initiation, maintenance and termination.

In summary, our single-locus live-cell measurements indicate that chromatin-mark–associated signals around endogenous genes can fluctuate reversibly within minutes and are coupled to transcriptional state switching. We propose that transcriptional bursts are embedded in a dynamically regulated chromatin environment shaped by co-recruited opposing enzymatic activities, creating feedback that can sustain bursts yet also promote timely termination. This temporal view helps reconcile stochastic bursting with robust gene regulation across developmental timescales (Fig. 6).

## Methods

### Cell lines and culture conditions

Wild-type mouse embryonic stem cells (mESCs; C57BL/6J, male) were used as the primary parental line in this study, and *Nanog*-, *Usp5*-, and *Dnmt3L*-STREAMING-tag knock-in lines ^11^ were derived from this background. A *Sox2* STREAMING-tag knock-in line ^11^ was generated independently using C57BL/6NCr (male) mESCs^10^. A complete list of cell lines and their strain backgrounds is provided in Supplementary Table 1. All mESC lines and their derivatives were cultured on 0.1% gelatin-coated plates at 37 °C in a humidified 5% CO₂ atmosphere. Naive pluripotency was maintained using 2i medium, which consisted of Dulbecco’s modified Eagle’s medium (DMEM) supplemented with 15% fetal bovine serum (FBS), 0.5 mM monothioglycerol, 1× MEM non-essential amino acids, 2 mM L-alanyl-L-glutamine, 1,000 U/mL leukemia inhibitory factor (LIF), 20 µg/mL gentamicin, 3 µM CHIR99021, and 1 µM PD0325901.

### Establishment of *Dnmt3L* STREAMING-tag knock-in mESCs

WT mESCs were transfected using Lipofectamine 3000 (Thermo Fisher Scientific) with pTV-Dnmt3L-STtag (2,000 ng), eSpCas9-EF-Dnmt3L (700 ng), and pKLV-PGKpuro2ABFP (Addgene plasmid #122372; 300 ng). As a control, cells were transfected with pTV-Dnmt3L-STtag (2,000 ng) and pKLV-PGKpuro2ABFP (300 ng) without the Cas9 plasmid (Supplementary Table 1). Twenty-four hours after transfection, cells were cultured in 2i medium supplemented with puromycin (1 µg/mL; FUJIFILM Wako Pure Chemical). One day after puromycin treatment, cells were passaged in their entirety onto gelatin-coated 10-cm dishes and cultured in 2i medium containing G418 (200 µg/mL; Nacalai Tesque). After 7 days of selection in 2i medium with 200 µg/mL G418, a stable G418-resistant cell population was obtained.

Twenty-four colonies were picked for further analysis. Genomic DNA was extracted and screened by genomic PCR to narrow down candidate clones. Candidate clones were then validated by Southern blotting as described previously ^44^. DIG-labelled probes were prepared using the PCR DIG Probe Synthesis Kit (Roche, Basel, Switzerland; 11636090910).

Cells established through this procedure are referred to as *Dnmt3L* STREAMING-tag knock-in cells in this study. To verify the edited locus, a genomic region spanning the target site was PCR-amplified from genomic DNA of the established cell line using primer-F (GATGTCTCAGGGCCCCTATTTCTT) and primer-R (TCGGTTCACTTTGACTTCGTACCT), and the amplicon was sequenced. This analysis revealed that the non-knock-in allele in this cell line carries a three-base deletion (Supplementary Fig. 1c).

### Establishment of cell lines stably expressing mTetR-YFP and MCP-CFP

Cell lines stably expressing fluorescently tagged mTetR and MCP were generated by piggyBac-mediated genomic integration. *Nanog*, *Sox2*, *Usp5*, and *Dnmt3L* STREAMING-tag knock-in cells ^11^ were seeded at 2.5 × 10⁵ cells per well in 24-well plates containing 0.5 mL 2i medium and allowed to attach for 1 h before transfection. For each well, cells were co-transfected with pCAG-hyPBase (50 ng), pLR5-CAG-TetR_W43F-3×mhYFP (75 ng), and pLR5-CAG-hMCP-3×mTurquoise2-NLS-2A-NLS-CLIP (375 ng) (Supplementary Table 1). Plasmids were delivered using Lipofectamine 3000 with P3000 reagent (Thermo Fisher Scientific) in reduced-serum Opti-MEM (Thermo Fisher Scientific) according to the manufacturer’s instructions. The medium was replaced with fresh 2i medium 24 h post-transfection and changed daily thereafter. On day 5, cells were expanded into 12-well plates.

On day 6, cells were labelled with CLIP-Cell TMR-Star (New England Biolabs) and subjected to fluorescence-activated cell sorting (FACS) using a BD FACSAria III cell sorter. A population co-expressing clearly high levels of monomeric hyperfolder YFP and CLIPtag was first enriched (P6; Supplementary Fig. 3a). Because CFP fluorescence could not be detected by our flow cytometer, CLIP fluorescence was used as a proxy for MCP-3×mTurquoise2 (CFP) expression (the two are co-expressed from the same transcript via a 2A peptide). After expansion, cells were re-labelled with CLIP-Cell TMR-Star and subjected to two additional rounds of FACS to isolate populations co-expressing moderate levels of YFP and CLIPtag (P7 and P10; Supplementary Fig. 3b,c). Multiple rounds were performed because cells expressing only YFP or only CLIPtag can occupy the broad intermediate gate, and repeated sorting increases the fraction of balanced double-positive cells. Eight days after the final sorting, individual colonies were manually picked and transferred to gelatin-coated 8-well chambered cover glasses. Clones exhibiting stable and moderate fluorescence levels were identified by fluorescence microscopy and used for downstream live-cell imaging experiments (Supplementary Table 1).

### Establishment of cells expressing SNAPtag fused protein of interest by lentivirus transduction

Lenti-X 293T cells (Clontech, 632180) were seeded at 1.0 × 10^6^ cells per well in 6-well plates (Thermo Fisher Scientific, 140675). After 24 h incubation, for each well, 7.5 µL Lipofectamine 2000 (Thermo Fisher Scientific, 11668019) was diluted in 250 µL Opti-MEM (Thermo Fisher Scientific, 31985062), gently mixed, and incubated for 5 min at room temperature. In parallel, a total of 3.0 µg plasmid DNA was prepared by mixing 1.5 µg lentiviral vector (e.g., pFUGW-Snap-mHdac1), 0.75 µg pMDLg/pRRE (Addgene plasmid #12251), 0.30 µg pRSV-Rev (Addgene plasmid #12253), and 0.45 µg pMD2.G (Addgene plasmid #12259), and the DNA mixture was diluted in 250 µL Opti-MEM (Supplementary Table 1). The diluted DNA and diluted Lipofectamine 2000 were then combined, gently mixed, and incubated for 20 min at room temperature to allow DNA–lipid complex formation. During complex formation, culture medium was replaced with 1.0 mL pre-warmed DMEM (Thermo Fisher Scientific, 11965-092) supplemented with 10% FBS. Subsequently, 500 µL of the DNA–lipid complexes was added to the cells. At 16 h post-transfection, the medium was replaced with 2.0 mL fresh DMEM supplemented with 10% FBS, and cells were cultured for an additional 48 h.

At 48 h post-transfection, lentivirus-containing supernatant was collected and clarified by filtration through a 0.45 µm filter (Millipore, SLHV033RS). The clarified supernatant was concentrated by adding one-third volume of Lenti-X Concentrator (Clontech, 631231) followed by gentle inversion and incubation at 4 ℃ for 30 min to overnight. Samples were centrifuged at 1,500 × g for 45 min at 4 ℃, and the supernatant was carefully removed without disturbing the pellet. Viral pellets were resuspended in 40 µL PBS per 6-well preparation, aliquoted (10 µL per tube), and stored at −80 ℃. *Nanog*, *Sox2*, *Usp5*, and *Dnmt3L* STREAMING-tag knockin cells mildly expressing mTetR-YFP and MCP-CFP mESCs, or *Nanog* or *Sox2* STREAMING-tag knockin cells mildly expressing mTetR-GFP and MCP-RFP (NSt-GR or SSt-GR)^11^ mESCs cultured under LIF + serum ES medium (Dulbecco’s modified Eagle’s medium (DMEM) supplemented with 15% FBS, 0.5 mM monothioglycerol, 1× MEM non-essential amino acids, 2 mM L-alanyl-L-glutamine, 1,000 U mL⁻¹ leukemia inhibitory factor (LIF), 20 µg mL⁻¹ gentamicin) were seeded at 5 × 10⁴ cells per well and exposed to lentiviral particles in the presence of 8 µg/ml polybrene (Sigma-Aldrich, H9268). Spinfection was performed by centrifugation at 1,000 × g for 2 h at 37 ℃, followed by incubation for an additional 30 min. The viral inoculum was then replaced with fresh 2i medium, and cells were cultured overnight. Cells were transferred to 12-well plates the following day and expanded under standard conditions. At 6 days post-infection, cells were labelled with JF646-SNAPtag ligand by incubation in 2i medium containing 300 nM ligand for 30 min. Cells were then washed three times with PBS and further incubated in fresh 2i medium for 30 min to allow removal of unbound ligand. Cells were dissociated into single-cell suspensions for fluorescence-activated cell sorting (FACS) using a BD FACSAria III cell sorter. For experiments using exogenously expressed SNAPtag fusions in the mTetR-YFP / MCP-CFP reporter background, a population expressing medium levels of SNAPtag was isolated (Supplementary Fig. 3d; Supplementary Table 1). For experiments using exogenously expressed SNAPtag fusions in the mTetR-GFP / MCP-RFP reporter background, we sorted cells into low/mid/high SNAPtag-expression bins (P4–P6; Supplementary Fig. 9a) and used the low-expression bin (P4) for subsequent imaging to minimize perturbation. For HDAC1/2/3-SNAPtag, immunoblotting confirmed that P4-sorted cells express the SNAPtag fusion at levels comparable to or below endogenous HDACs (Supplementary Fig. 9b; Supplementary Table 1).

### Establishment of cells expressing SNAPtag fused protein of interest by transposon-mediated integration

*Nanog* or *Sox2* STREAMING-tag knockin cells mildly expressing mTetR-GFP and MCP-RFP (NSt-GR or SSt-GR)^11^ mESCs (2.5 × 10^5^) were plated into each well of a 24-well plate, and after 1 h, the following transfection reagents were mixed: 50 ng pCAG hyPBase, 75 ng pLR5-CAG-TetR_W43F-3xmNG, 275 ng pLR5-CAG-hMCP-mScarlet-I-NLS, and 100 ng of SNAPtag expression vector (e.g., pLR5-CAG-SNAPtag-mEP300; Supplementary Table 1). To each of these, 25 μL of reduced serum Opti-MEM and 1 μL of P3000 reagent were added. In a separate tube, 25 μL of Opti-MEM reduced serum medium and 1.8 μL of Lipofectamine 3000 were added per reaction and mixed well. The P3000 and Lipofectamine 3000 media were mixed in equal volumes and incubated at RT for 15 min. After 24 h (day 2), the medium was replaced with 2i medium. After another 24 h (day 3), the medium was replaced with 2i medium, and the cells were passaged in 12-well plates on day 5. On day 6, the cells were incubated in 2i medium containing 300 nM SNAP-Cell 647-SiR (New England Biolabs, Ipswich, MA, USA, S9102S) for 30 min at 37 °C and 5% CO_2_. The cells were washed three times with 2i medium and incubated at 37 °C and 5% CO_2_ for another 30 min. The cells were collected following treatment with trypsin. For experiments using exogenously expressed SNAPtag fusions in the mTetR-YFP / MCP-CFP reporter background, a population expressing medium levels of SNAPtag was isolated (Supplementary Fig. 3d; Supplementary Table 1). For experiments using exogenously expressed SNAPtag fusions in the mTetR-GFP / MCP-RFP reporter background, we sorted cells into low SNAPtag-expression bins (P4; Supplementary Fig. 9a) and used the low-expression bin (P4) (Supplementary Fig. 9a; Supplementary Table 1). Sorted cells were seeded into gelatin-coated 6 cm dishes.

### CRISPR-Cas9-mediated SNAPtag knock-in

*Nanog* or *Sox2* STREAMING-tag knockin cells mildly expressing mTetR-GFP and MCP-RFP (NSt-GR and SSt-GR cells^11^) (1.25 × 10^5^) were plated into each well of a 24-well plate, and after 1 h, the transfection reagents were mixed. In a tube, 500 ng of targeting vector (e.g., pTV-FLAG-SNAPtag-Chd4), 250 ng of CRISPR vector (e.g., eSpCas9-EF-Chd4), and 75 ng of pKLV-PGKpuro2ABFP (Addgene plasmid #122372) were mixed (Supplementary Table 1). To each of these, 31 μL of reduced serum Opti-MEM and1.25 μL of P3000 reagent were added. In a separate tube, 31 μL of reduced serum Opti-MEM and 2.25 μL of Lipofectamine 3000 were added per reaction and mixed well. The P3000 and Lipofectamine 3000 media were mixed in equal volumes and incubated at RT for15 min. After 24 h (day 2), the cells were treated with 1 μg/mL puromycin in 2i medium. The medium was replaced with fresh 2i medium after another 24 h (day 3). Every 24 h, the medium was replaced with fresh 2i medium. On day 6, the cells were incubated for 30 min in 2i medium containing 300 nM SNAP-Cell 647-SiR at 37 ℃ and 5% CO_2_.The cells were washed three times with 2i medium and incubated at 37 ℃ and 5% CO_2_ for another 30 min. The cells were collected by trypsin treatment, and SNAPtag signal-positive cells were sorted using a BD FACSAria III cell sorter and seeded into gelatin-coated 6 cm dishes. The medium was changed once every 2 days. Twenty-four colonies were picked on day 8 after FACS. Genomic DNA extracted from these cells was used for genomic PCR to narrow down the candidate cell lines. Candidate clones were further analyzed using Southern blotting. Probes were prepared using a PCR DIG Probe Synthesis Kit (Roche Diagnostics, Mannheim, Germany). The SNAPtag was knocked into *Chd4* and *Sin3a* for NSt-GR and SSt-GR mESCs, and *Sox2* only for NSt-GR mESCs, as described above. See Supplementary Table 1 for the plasmids used in this study. We confirmed that these SNAPtag knock-in cell lines have a growth rate comparable to that of the parental cell lines, suggesting that the effect of SNAPtag knock-in on the cells is negligible.

### Snapshot imaging of SNAPtag-expressing cells

*Dnmt3L* expression is known to be reduced in mESCs maintained in 2i medium compared with cells cultured in LIF + serum ES medium. To facilitate robust detection of *Dnmt3L* transcription, *Dnmt3L* STREAMING-tag knock-in-derived cells were pre-adapted to LIF + serum ES medium for three passages before the experiments described below. All other cell lines were maintained in 2i medium prior to imaging.

SNAPtag-expressing mESCs were seeded onto laminin-511 (Biolamina)-coated 8-well glass-bottom (Cellvis, C8-1.5H-N) chamber slides at 5 × 10⁴ cells per well and cultured overnight in their respective maintenance media (2i medium for all lines except *Dnmt3L* STREAMING-tag knock-in-derived cells, which were maintained in LIF + serum ES medium).

For SNAPtag labeling, cells were incubated with JF646-SNAPtag ligand in the corresponding FluoroBrite-based imaging medium (300 nM for 30 min at 37 °C, unless noted otherwise). FluoroBrite-based 2i medium consisted of FluoroBrite DMEM (Thermo Fisher; A1896701) supplemented with 15% FBS, 0.5 mM monothioglycerol, 1× MEM non-essential amino acids, 2 mM L-alanyl-L-glutamine, 1,000 U/mL LIF, 20 µg/mL gentamicin, 3 µM CHIR99021, 1 µM PD0325901, and 1:1,000 VectaCell Trolox Antifade Reagent for Live Cell Imaging (Vector Laboratories, CB-1000). FluoroBrite-based LIF + serum ES medium consisted of FluoroBrite DMEM supplemented with 15% FBS, 0.5 mM monothioglycerol, 1× MEM non-essential amino acids, 2 mM L-alanyl-L-glutamine, 1,000 U/mL LIF, 20 µg/mL gentamicin, and 1:1,000 VectaCell Trolox Antifade Reagent.

Cells were washed three times with the corresponding FluoroBrite-based medium to remove unbound ligand and then incubated in fresh FluoroBrite-based medium for an additional 30 min to allow ligand clearance. Immediately before imaging, the medium was replaced with fresh FluoroBrite-based medium. In some experiments, to minimize handling, cells were alternatively labeled by incubation in FluoroBrite-based imaging medium containing 100 nM JF646-SNAPtag ligand for ≥90 min at 37 °C and imaged directly without washing or medium exchange.

Live-cell fluorescence imaging was performed on a Nikon Ti-2 microscope equipped with a CSU-W1 spinning-disk confocal unit (Yokogawa), a 100× Nikon Plan Apo λ oil-immersion objective (NA 1.4), and an iXon Ultra EMCCD camera (Andor), controlled by NIS-Elements (v5.11.01; Nikon). Cells were maintained in a stage-top incubator at 37 ℃ with 5% CO₂. The microscope was equipped with 405-, 445-, 488-, 515-, 561-, and 637-nm lasers (LightHUB ULTRA, Omicron) and an ASI MS-2000 piezo stage (Applied Scientific Instrumentation). Z-stack images spanning 5 µm were acquired at 500-nm steps (11 optical sections; 130 nm per pixel).

For cells expressing MCP-CFP and mTetR-YFP, imaging was performed using three channels corresponding to mTetR-YFP (515-nm excitation), MCP-CFP (445-nm excitation), and SNAPtag (637-nm excitation). For cells expressing MCP-RFP and mTetR-GFP, imaging was performed using three channels corresponding to mTetR-GFP (488-nm excitation), MCP-RFP (561-nm excitation), and SNAPtag (637-nm excitation).

### Snapshot image processing and analysis of STREAMING-tag cells

All snapshot STREAMING-tag image analyses were performed using custom Python scripts with Cellpose^45^, Big-FISH^46^, and Trackpy. Nuclei were first segmented using Cellpose (nuclei model; diameter set to 100 px), and segmentation masks were visually inspected.

To identify the tagged genomic locus, mTetR foci were detected per nucleus using Big-FISH (detection.detect_spots) in the 3D mTetR channel within the nuclear bounding box, using an adaptive threshold defined as one twentieth of the mean mTetR intensity measured for that nucleus; among candidate spots, the brightest spot within the nuclear mask was selected as the locus and its (z, y, x) coordinates were recorded. For downstream quantification, a locus-centered crop was extracted at the z-plane corresponding to the selected mTetR spot: a 19 × 19 pixel region was cut from all three channels (SNAP/factor, MCP, and mTetR). Pixel size was 130 nm/pixel.

Relative enrichment metrics were computed from locus-centered crops using Trackpy-based center-of-mass intensity estimation. Briefly, for a given coordinate, a “core” intensity was calculated from refine_com as (mass/area), where area = pad²·π with pad = 3 pixels, and normalized by the mean intensity of the entire 19 × 19 crop to yield a relative ratio *r* = *I_core_*/*I_mean_*. The locus quality metric *r_mTetR_* was computed at the crop center; cells with *r_mTetR_* below threshold were excluded. The transcription metric *r_mcp_* was computed for the brightest MCP focus detected in the crop (Trackpy locate followed by 2D Gaussian refinement). A locus was classified as transcriptionally Active state when an MCP focus satisfied (i) *r_mcp_* above threshold and (ii) spatial proximity to the locus center within 3 pixels (390 nm); otherwise it was classified as Inactive state. For all analyses used in this study, thresholds were unified to *r_mTetR_* = 1.30 and *r_mcp_* = 1.15.

To visualize factor enrichment around the locus, locus-centered images were grouped by transcriptional state (Active vs Inactive) and pixel-wise mean images were computed for each channel. Random nuclear control crops (sampled away from the locus and processed identically) were used to estimate baseline background; the mean random image was subtracted prior to visualization and radial profiling. Radial intensity profiles were computed for each locus instance by averaging the (background-subtracted) intensity within concentric 1-pixel annuli around the crop center, and then summarized as the mean ± s.e.m. across locus instances for each state. For statistical testing, Active and Inactive annulus-intensity distributions were compared at each radius using a two-sided Mann-Whitney U test, followed by Benjamini-Hochberg correction across radii (*q* < 0.05). For visualization, background-subtracted images were linearly scaled within each marker to a common display range (arbitrary units) using the same scaling factor for Active and Inactive conditions.

For variability and correlation analyses, SNAP/factor signal at the locus center was quantified as the mean intensity within a 3 × 3 pixel window centered on [9,9]. Distributions of center SNAP intensity were visualized as violin plots after log_10_ transformation and compared between Active and Inactive cells using a two-sided Mann-Whitney U test. To assess coupling between SNAP accumulation and transcriptional activity, a center-based MCP relative metric (*r_mcp,center_*) was computed at the locus center using the same core/mean normalization (pad = 3), and Spearman correlation coefficients between center SNAP intensity and (*r_mcp,center_*) were calculated across cells and summarized as heatmaps.

### Genome-wide enrichment analysis of ChIP-seq/ATAC-seq bigWig tracks at loci analyzed by STREAMING-tag

Publicly available ChIP-seq and ATAC-seq bigWig tracks were analyzed on the mouse genome (mm10) to assess the genome-wide context of chromatin marks and transcription-associated factors at loci analyzed by STREAMING-tag. The mm10 genome (autosomes and sex chromosomes) was partitioned into non-overlapping fixed bins of 0.5 Mb. For each track, mean signal per bin was computed from the bigWig file using pyBigWig (stats, type = “mean”), generating a genome-wide background distribution of bin-wise signals.

For genes of interest (*Nanog*, *Sox2*, *Usp5* and *Dnmt3L*), transcription start sites (TSSs) were inferred from the annotated strand (TSS = gene start for ‘+’ strand and gene end for ‘-’ strand). A 0.5-Mb TSS-centered window was then defined for each gene; when a window crossed a chromosome boundary, it was shifted to remain within the chromosome while preserving window length. For each track, the mean bigWig signal was extracted for each gene window.

To enable comparison across targets with different signal scales, we computed per-target joint z-scores. For each target *t*, genome-wide bin values and the four gene-window values were combined into a single distribution, and the mean (*μ_t_*) and standard deviation (*σ_t_*) were computed from the combined values. Each value *x* was then standardized as *z_joint_* = (*x* - *μ_t_*)/*σ_t_*. For visualization (Supplementary Fig. 5e), boxplots show the distribution of *z_joint_* across genome-wide bins for each target, and gene-window *z_joint_* values were overlaid as colored points.

### Reanalysis of published seq-DNA/RNA/IF-FISH datasets

We reanalyzed publicly available seq-DNA/RNA/IF-FISH datasets reported by Ohishi et al.^25^. The seq-DNA-FISH coordinate table provides, for each detected allele-locus instance, the field of view (FOV), cell/allele identifiers (e.g., IntraFOV_Cell_ID, Cell_ID, and Allele_lID), the locus identifier (Locus ID), and estimated 3D coordinates (x_nm, y_nm, z_nm). Corresponding 3D immunofluorescence (IF) image stacks were used to quantify local IF signal at each seq-DNA-FISH locus.

To quantify IF signal at the seq-DNA-FISH loci, we extracted voxel-wise fluorescence intensities from 3D TIFF stacks for each IF marker (CHD4, BRD4, KDM1A, OTX2, MED12, SIN3A, SOX2, TCF3, ESRRB, and H3K27ac). For each locus, the reported nanometre-scale coordinates (x_nm, y_nm, z_nm) were converted to integer voxel indices by dividing by the voxel size (130 nm in x, y and z) and taking the floor of each coordinate (x_int = floor(x_nm / 130), y_int = floor(y_nm / 130), z_int = floor(z_nm / 130)). TIFF stacks were loaded using tifffile in Python and indexed as (z, y, x) arrays; the raw intensity at (z_int, y_int, x_int) was recorded as the IF signal for that marker at the locus.

Transcriptional activity states were assigned using the seq-RNA-FISH table provided by Ohishi et al., which enumerates only transcriptionally active (“ON”) allele-gene instances. A unique allele key was defined by concatenating FOV, cell, and allele identifiers to avoid ambiguity across fields of view. Each DNA/IF allele-locus instance was labeled Active if an identical entry was present in the seq-RNA-FISH table, and Inactive otherwise.

To compare IF enrichments across loci while accounting for gene-specific intensity distributions, IF signals were standardized using gene-wise z-scores. For each gene and each IF marker, the mean and standard deviation were computed across all alleles of that gene irrespective of transcriptional state. Z-scores were calculated as

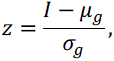

where *I* is the raw IF intensity for an allele, and *µ_g_* and *σ_g_* are the gene-specific mean and standard deviation for the marker. Cases with *σ_g_* = 0 were set to *z* = 0.

For each gene and IF marker, we computed: Δ*z* = median(*z|Active*) − median(*z|Inactive*). Genes were retained if at least one Active allele was observed (Active alleles >0%) and both Active and Inactive instances were available for Δ*z* estimation. Genes passing these criteria (67 genes) were ranked by Active allele ratio and stratified into quartiles for downstream analyses (Fig. 2b and Supplementary Fig. 6a).

For each IF marker, we summarized gene-wise state bias by computing Δ*z* values for each gene, defined as the difference between the median gene-wise z-score in transcriptionally Active alleles and that in Inactive alleles. Genes were ranked by their Active-allele ratio and grouped into quartiles. For each marker and quartile, the distribution of per-gene Δ*z* values was visualized using boxplots with individual gene values overlaid. To test whether Δ*z* values were systematically shifted from zero, we applied two-sided Wilcoxon signed-rank tests to the set of per-gene Δ*z* values. *P* values were adjusted for multiple testing using the Benjamini–Hochberg procedure to control the false-discovery rate (BH-FDR), and adjusted *q* values were used for significance annotation (stars). The star thresholds were: one star for *q* ≤ 0.05, two stars for *q* ≤ 0.01, three stars for *q* ≤ 0.001, and four stars for *q* ≤ 0.0001.

### Re-analysis of published sci-mtChIL-seq datasets

Published sci-mtChIL-seq data generated in NIH3T3 cells (RNAPII Ser2 phosphorylation paired with H3K27ac, H3K4me3, and H3K27me3) were re-analyzed using processed gene × cell count matrices ^28^. For each RNAPII-target pair, unique fragments were summarized at the gene level by counting RNAPII fragments within a promoter-proximal window around the transcription start site (TSS; −1 kb to +5 kb) and counting paired-target fragments within a symmetric TSS-centered window (TSS ±5 kb). Paired-target counts were converted to counts-per-thousand (CPT) per cell.

Genes were restricted to those with detectable expression in bulk RNA-seq, and the top 5,000 genes with the highest RNAPII counts were retained for downstream analyses. For each gene, cells were classified as transcriptionally Active state if the RNAPII count was >0 and Inactive otherwise. For each (target, gene) record, the mean paired-target CPT in Active and Inactive cells and the numbers of Active/Inactive (*N_Active_* and *N_Inactive_*, respectively) cells were computed.

To rank genes by activity frequency, we computed, for each gene, the fraction of Active cells,

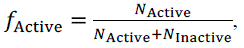

and ranked genes in descending order of *f_Active_*. To assess how coupling depends on transcriptional activity frequency, genes in the RNAPII-covered set were ranked by *f_Active_* and partitioned into consecutive 500-gene bins (ranks 1–500 to 4501–5000), and downstream comparisons were performed separately within each bin. For each gene, we computed the ratio between the mean paired-target CPT in Active cells and that in Inactive cells, after adding a pseudocount of 0.01 to both means to avoid division by zero. Deviations from a median ratio of 1 were assessed using a two-sided one-sample Wilcoxon signed-rank test on log_2_-transformed ratios, and *P* values were corrected for multiple testing using the Benjamini-Hochberg procedure.

### Chromatin state annotation and chromatin-state–specific enrichment analysis

Public ChIP-seq and ATAC-seq datasets in mouse embryonic stem cells (mm10) were obtained from ENCODE and the NCBI Sequence Read Archive (SRA) (accessions and file identifiers are listed in Supplementary Table 2). For chromatin state model learning, histone modification and CTCF datasets available as BAM files were downloaded directly from ENCODE. For datasets available only as SRA files (e.g., H3K4me2 and ATAC-seq), SRA files were converted to FASTQ using SRA Toolkit (fasterq-dump with split output for paired-end reads). Reads were filtered using fastp (quality threshold Phred 20; minimum read length 30; automatic adapter detection), aligned to the mm10 reference genome using Bowtie2 (paired-end alignment for H3K4me2; single-end alignment for the ATAC-seq dataset as provided), and converted to coordinate-sorted and indexed BAM files using samtools. A chromosome size file for mm10 was generated from the reference FASTA index (samtools faidx) for downstream ChromHMM processing.

Genome-wide chromatin state segmentation was performed using ChromHMM (v1.25)^47^. ChIP-seq/ATAC-seq signals were binarized at 200-bp resolution using ChromHMM BinarizeBam with default settings. A 12-state Hidden Markov Model was learned using ChromHMM LearnModel (bin size 200 bp; parallel computation enabled). Chromatin states were assigned biologically interpretable labels based on their emission patterns across marks and known functional associations. The final state labels used in this study were: Heterochromatin, Intergenic region, Weakly repressed Polycomb, Repressed Polycomb, Bivalent promoter, Weak enhancer, Strong enhancer, Enhancer, Transcription transition, Transcription elongation, Insulator, and Active promoter. The annotated 12-state segmentation was exported as BED files and used as a fixed genomic partition for downstream enrichment analyses. ChromHMM emission probabilities shown in the heatmap correspond to the probability of observing each mark given a chromatin state.

To quantify how transcription-associated factors and HDAC-related assemblies distribute across ChromHMM-defined chromatin states, genome-wide signal tracks (bigWig) for HDACs and associated cofactors (including HDAC1/2/3/8, CHD4, SIN3A, NCOR1/NCOR2 and SIRT6), and for activation-associated and transcription-related factors (including EP300, BRD4, RNAPII and RNAPII CTD phosphorylation marks) were obtained from ENCODE and/or ChIP-Atlas (track sources listed in Supplementary Table 2). For each factor and each chromatin state, mean signal was computed by aggregating bigWig signal values across all genomic intervals assigned to that state and normalizing by the total base-pair coverage of the state, yielding a per-base average signal for each factor–state pair. To facilitate comparison of relative state preferences across factors with different dynamic ranges, the resulting factor-by-state matrix was standardized by computing row-wise Z-scores across states for each factor. ChromHMM emission probabilities (state-by-mark) and the Z-scored factor enrichment matrix (state-by-factor) were visualized as aligned heatmaps.

### Re-analysis of MARCS data

Precomputed MARCS PTM-response results (Estimated effects for chromatin features (proteins) and Estimated effects for chromatin features (curated complexes)) were downloaded from the MARCS Readers portal (Helmholtz Munich; accessed on https://marcs.helmholtz-munich.de/downloads). The effect estimates reported by MARCS (PTM contribution to the H/L ratio on a log_2_ scale) and the accompanying multiple-testing-adjusted significance values provided in the workbooks were used for plotting without re-estimation. For the three chromatin features H3ac, H4ac, and H3K4me3, effect estimates for selected HDAC-associated factors (HDAC1, HDAC2, HDAC3, NCOR1, NCOR2, SAP30, and SIN3A) were extracted from the protein-level workbook. When multiple effect entries were available for a given gene and mark, the median effect was used. The resulting effect values were plotted as a dot plot across chromatin features (x-axis: effect in log_2_ H/L; y-axis: chromatin feature).

For curated complexes, volcano plots were generated separately for H3ac, H4ac, and H3K4me3 using the complex-level workbook (x-axis: reported median effect; y-axis: −log_10_(FDR)). Complexes were categorized and color-coded as “Up” (effect ≥ 0.5 and FDR ≤ 0.01), “Down” (effect ≤ −0.5 and FDR ≤ 0.01), or “significant with small effect” (|effect| < 0.5 and FDR ≤ 0.01). Complexes containing HDAC subunits were identified by the presence of the substring “HDAC” in the workbook field listing measurable subunits (e.g., “Proteins for which response was measurable”), highlighted with a distinct marker, and annotated by complex name.

### Western blotting

Cells were rinsed twice with phosphate-buffered saline (PBS, NacalaiTesque, 14249-24), detached with trypsin, and pelleted by centrifugation at 190× g for 2 min at 20 °C. After counting, cells were washed twice again with PBS and lysed in lysis buffer (0.5% Triton X-100 (Sigma-Aldrich, T8787-100ML), 150 mM NaCl (Wako, 191-01665), and 20 mM Tris-HCl [pH 7.5]) at a concentration of 2 × 10^6^ cells per 100 μL. Lysates were heated at 95 °C for 5 min and clarified using a QIAshredder homogenizer (Qiagen, 79656). Proteins were separated by 5–20% gradient SDS-PAGE. Prior to transfer, gels were briefly equilibrated in 20% ethanol for 5–10 min and rinsed in water. Proteins were transferred to PVDF membranes using an iBlot® 2 Dry Blotting System with iBlot® 2 Transfer Stacks (PVDF, mini; program P0) (Thermo Fisher Scientific).

Membranes were immediately immersed in TBST (TBS containing 0.1% Tween-20) and blocked in 0.3% (w/v) skim milk in TBST for at least 1 h at room temperature with gentle agitation. Membranes were incubated with primary antibodies diluted in blocking buffer overnight at 4 °C, washed with TBST (15 min once and 5 min three times), and incubated with appropriate HRP-conjugated secondary antibodies diluted in blocking buffer for 90 min at room temperature. After washing with TBST (15 min once and 5 min four times), chemiluminescence was developed using SuperSignal™ West Dura Extended Duration Substrate (Thermo Fisher Scientific) according to the manufacturer’s instructions and imaged on a CCD-based chemiluminescence imager. The primary antibodies used were anti-GAPDH (1:5000; Cell Signaling Technology, 5174, RRID:AB_10622025), anti-HDAC1 (1:1000; GeneTex, GTX100513, RRID:AB_1240929), anti-HDAC2 (1:1000; Cell Signaling Technology, 5113T, RRID:AB_10624871), and anti-HDAC3 (1:2000; GeneTex, GTX113303, RRID:AB_10721050).

### Time-lapse imaging of SNAPtag-expressing cells

SNAPtag-expressing mESCs were seeded onto laminin-511 (Biolamina)-coated 8-well glass-bottom (Cellvis) chamber slides at 5 × 10⁴ cells per well and cultured overnight in 2i media.

For SNAPtag labeling, cells were incubated with 300 nM JF646-SNAPtag ligand for 30 min at 37 ℃ in the 2i containing FluoroBrite-based imaging medium. Cells were washed three times with the 2i containing FluoroBrite-based medium to remove unbound ligand and then incubated in fresh 2i containing FluoroBrite-based medium for an additional 30 min to allow ligand clearance. Immediately before imaging, the medium was replaced with fresh 2i containing FluoroBrite-based medium containing 100 nM JF646-SNAPtag ligand.

Live-cell fluorescence imaging was performed on a Nikon Ti-2 microscope equipped with a CSU-W1 spinning-disk confocal unit (Yokogawa), a 100× Nikon Plan Apo λ oil-immersion objective (NA 1.4), and an iXon Ultra EMCCD camera (Andor), controlled by NIS-Elements (v5.11.01; Nikon). Cells were maintained in a stage-top incubator at 37 ℃ with 5% CO₂. The microscope was equipped with 405-, 445-, 488-, 515-, 561-, and 637-nm lasers (LightHUB ULTRA, Omicron) and an ASI MS-2000 piezo stage (Applied Scientific Instrumentation). Z-stack images spanning 6 µm were acquired at 300-nm steps (21 optical sections; 130 nm per pixel).

Imaging was performed using three channels corresponding to mTetR-GFP (488-nm excitation), MCP-RFP (561-nm excitation), and SNAPtag (637-nm excitation). Time-lapse images were acquired at 2-min intervals for a total duration of 2 h.

### Time-lapse image preprocessing and analysis

Time-lapse movies were acquired at 2-min intervals for 2 h. Raw ND2 files were imported into Fiji using Bio-Formats and exported as TIFF hyperstacks. Each fluorescence channel was denoised using a 3D median filter (radius 1 pixel/voxel in x, y, and z). To enhance locus contrast, diffuse background in the mTetR channel was suppressed by subtracting a 3D Gaussian-blurred copy (σ = 5 pixels). Maximum-intensity projections (MIPs) were generated for visualization and segmentation.

Single-cell segmentation and tracking were performed on 2D maximum-intensity projections (MIPs). When the SNAPtag MIP provided stable masks across frames, the SNAPtag channel was used for segmentation; otherwise, a composite segmentation image was generated by robustly normalizing the MCP and mTetR MIPs separately (1st–99.8th intensity percentiles) and summing the two normalized images to improve cell-to-background contrast. The selected segmentation image was then smoothed (Gaussian σ = 1) and segmented by Otsu thresholding. Masks were refined by morphological opening (radius 2) and closing (radius 3), hole filling, and removal of small objects (<1500 pixels). A single cell was tracked over time by linking mask centroids with a nearest-neighbor rule (maximum displacement 60 pixels), and movies were cropped to retain the tracked cell throughout the time series.

Gene-locus positions (mTetR) and nascent RNA foci (MCP) were detected in 3D using Big-FISH with voxel sizes of 130 nm (xy) and 300 nm (z) and theoretical spot radii of 180 nm (xy) and 400 nm (z). Candidate mTetR spots were linked across frames by selecting the nearest detection to the previous-frame coordinate (maximum distance 30 pixels). Spot tracks were manually curated in napari and refined computationally by center-of-mass localization within a local 3D neighborhood (7×7 pixels in xy and 3 z-slices) after local background subtraction. For each frame, the cell mask was additionally constrained by selecting the connected component containing the refined mTetR coordinate to ensure locus assignment to the tracked cell.

Locus-centered intensity traces were quantified from the SNAPtag and MCP channels as the mean intensity within a 3D window centered on the refined mTetR coordinate (±3 pixels in xy and ±2 z-slices). Transcriptional activity was called per frame by searching for an MCP focus within 3 pixels of the mTetR position and requiring a signal-to-background ratio >1.2 relative to local background measured in the same neighborhood.

Photobleaching correction was applied to the SNAPtag locus-centered trace by computing a reference mean-intensity series from a nucleus-like region within the tracked cell: a nucleus mask was obtained by Otsu thresholding of the SNAPtag MIP within the cell mask and selecting the largest connected component (fallback: the full cell mask). An exponential decay model was fit to the reference series, enforcing decay-only behavior, and the resulting multiplicative correction factor was capped at 3.0. The photobleaching-corrected locus-centered traces were stored as “detrended” columns (e.g., ch1_detrended and ch2_detrended). MCP bleaching correction was applied only when clear decay was detected in the reference trace (auto mode based on a head-to-tail intensity ratio criterion).

For correlation analyses, time series were reindexed to a continuous frame axis (expected 60 frames) without interpolation; missing frames were kept as NaN. Signals were z-scored within each cell, and cross-correlations were computed for lags up to ±10 frames using only overlapping finite pairs (minimum 3 pairs per lag), retaining the number of contributing pairs as weights. Group-mean correlation functions were computed as weighted averages across cells and baseline-corrected by subtracting the weighted mean correlation over |lag| = 7–10 frames. Uncertainty bands were estimated by cell-wise bootstrap resampling (3,000 replicates; 95% percentile intervals). For visualization, correlation curves were optionally smoothed by variable-width lag binning (multitau scheme).

Lead–lag asymmetry was quantified using a lead–lag index (LLI) computed per cell from the baseline-corrected cross-correlation as:

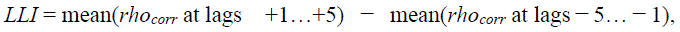

where negative lag corresponds to the SNAPtag/factor signal leading MCP under our convention. Thus, positive LLI values indicate that MCP tends to lead, whereas negative LLI values indicate that the SNAPtag/factor signal tends to lead. Statistical significance was assessed using within-cell circular time-shift permutation tests (1,000 permutations), generating a null distribution of the median LLI across cells. Two-sided permutation p-values were corrected for multiple comparisons using the Benjamini–Hochberg procedure.

### Time-lapse imaging of cells treated by HDAC inhibitors

NSt-GR and SSt-GR cells^11^ were seeded onto laminin-511 (Biolamina)-coated 8-well glass-bottom (Cellvis) chamber slides at 5 × 10⁴ cells per well and cultured overnight in their respective 2i media.

Immediately before imaging, the medium was replaced with fresh 2i containing FluoroBrite-based medium. HDAC inhibitors (2 µM TSA (Wako Chemicals, 203-17561) or 10 µM RGFP966 (Selleck Chemicals, S7229)) or vehicle control (DMSO) were then added, and live-cell fluorescence imaging was initiated on a Nikon Ti-2 microscope equipped with a CSU-W1 spinning-disk confocal unit (Yokogawa), a 100× Nikon Plan Apo λ oil-immersion objective (NA 1.4), and an iXon Ultra EMCCD camera (Andor), controlled by NIS-Elements (v5.11.01; Nikon). Cells were maintained in a stage-top incubator at 37 ℃ with 5% CO₂. The microscope was equipped with 405-, 445-, 488-, 515-, 561-, and 637-nm lasers (LightHUB ULTRA, Omicron) and an ASI MS-2000 piezo stage (Applied Scientific Instrumentation). Z-stack images spanning 6 µm were acquired at 300-nm steps (21 optical sections; 130 nm per pixel). Imaging was performed using three channels corresponding to mTetR-GFP (488-nm excitation), MCP-RFP (561-nm excitation), and SNAPtag (637-nm excitation). Time-lapse images were acquired at 1-min intervals for a total duration of 1 h.

### Analysis of HDAC inhibitor time-lapse data

Time-lapse datasets were analyzed using a custom Python pipeline that takes two-channel TIFF movies (time × channel × y × x) generated from per-frame 2D maximum-intensity projections (MIPs) of the acquired z-stacks. The input stacks contained the MCP channel (nascent transcription reporter) and the mTetR channel (locus marker). Frames were analyzed at Δ*t* = 1 min; pixel size was set to 130 nm.

For each frame, each channel was robustly normalized using the median and median absolute deviation (MAD) to facilitate spot detection. Candidate spots were detected on normalized images using Big-FISH spot detection with channel-specific normalized thresholds (THRESH_NORM = 1.25 for MCP and 1.10 for mTetR). When cell label masks were available, candidate spots were assigned to individual cells by the label value at the spot coordinate; otherwise, the field of view was treated as a single cell region.

To obtain a single locus trajectory per cell, at most one representative spot per cell per frame was selected for each channel and linked across consecutive frames by nearest-neighbor association with a maximum displacement of 3 pixels (390 nm). Tracks shorter than three consecutive frames were excluded. Transcriptional state was binarized per frame based on MCP spot detection at the locus. Specifically, a frame was classified as Active state when an MCP spot was detected within 3 pixels (390 nm) of the locus position defined by the mTetR trajectory. When an mTetR spot was not detected at the standard threshold in a given frame, we first attempted a local rescue by searching for a local intensity maximum within 6 pixels of the expected locus position using a lowered normalized threshold (0.935; 85% of the standard mTetR threshold). If no spot was recovered, the locus position was inferred from nearest-neighbor linking across adjacent frames, and the same proximity criterion was applied. To suppress spurious single-frame activations, isolated Active calls without temporal support from adjacent frames were relabeled as Inactive state.

From the resulting binary time series, contiguous Active and Inactive runs were extracted for each cell, and run duration was recorded in minutes. Runs overlapping the first or last frame of a movie were treated as left- or right-censored, respectively, and censoring annotations were retained. Frame-level outputs (Active/Inactive calls) and run-level outputs (run duration and censoring) were aggregated across movies and cells for each condition and locus.

To compare inhibitor conditions, we generated (i) representative state rasters, (ii) cell-level duty cycles (fraction of frames classified as Active per cell), and (iii) dwell-time distributions. Dwell-time distributions were analyzed using Kaplan-Meier survival curves, treating right-censored runs as censored observations and excluding left-censored runs in the primary analysis.

## Acknowledgments

We thank Dr. Luke D. Lavis (Janelia Research Campus, HHMI) for providing the JF646-SNAPtag ligand. We also thank Dr. Tuncay Baubec (Utrecht University) for kindly providing the H3K4me3 ChromID plasmid. This work was partly performed in the Cooperative Research Project Program of the Medical Institute of Bioregulation, Kyushu University. This work was supported by the following research grants: Japan Society for the Promotion of Science (JSPS) JP24H02326 (to H.Oc.), JP24H02323 (to Y.O.), JP23H00372 (to Y.O.), JP24H02325 (to H.K.); JST CREST Program JPMJCR23N3 (to H.Oc.); AMED BINDS JP22ama121017j0001 (to Y.O.); AMED ASPIRE 27jf0126008h0002 (to Y.O.); the Takeda Science Foundation to H.Oc; Medical Research Center Initiative for High Depth Omics, Kyushu University (to H.Oc. and Y.O.); MEXT Promotion of Development of a Joint Usage/Research System Project: Cooperative Research Project Program (to H.Oc. and Y.O.); MEXT Promotion of Development of a Joint Usage/Research System Project: Coalition of Universities for Research Excellence Program (CURE) JPMXP1323015486 (to H.Oc. and Y.O.); Medical Research Center Initiative for High Depth Omics (to H.Oc. and Y.O.); and NIG-JOINT 15R2025 and 42A2025 (to H.Oc.).

## Author contributions

H.Oc. conceived, designed, and supervised the study.

H.Oc. and Y.D. established all the knock-in cells.

H.Oc., X.G., C.K., S.U., Y. K., H.Oh. established mintbody-SNAPtag and SNAP-tagged protein expressing cells.

T.F., A.H., Y.O., and H.Oc. performed sci-mtChIL-seq re-analysis.

X.G., C.K., and H.Oc. performed STREAMING-tag and SNAPtag imaging experiments and analyzed the data.

H.Oc, X.G., C.K., and H.K. interpreted the data and wrote the manuscript.

## Additional information

### Materials & Correspondence

Supplementary Information is available for this paper.

Correspondence and requests for materials should be addressed to Hiroshi Ochiai.

## Statistical information

Statistical analyses were performed using custom scripts in Python. Unless otherwise stated, all hypothesis tests were two-sided and non-parametric tests were used because normality was not assumed. Imaging experiments were independently repeated twice with similar qualitative trends. Because absolute fluorescence intensities can vary modestly between independent imaging sessions, primary quantification was performed using a single replicate as indicated for each figure, and the second replicate was used as an independent confirmation of reproducibility.

Definitions of n and replicates: For snapshot imaging analyses, n denotes individual locus instances (one locus per nucleus/cell) that passed the predefined quality filters described in Methods. For time-lapse analyses, n denotes individual cells for duty-cycle analyses and individual Active/Inactive runs for dwell-time analyses (Kaplan-Meier), as indicated in each panel. For re-analyses of published datasets, n denotes genes (or allele instances where applicable).

Error bars and summary statistics: Error bars represent the standard error of the mean (s.e.m.) unless otherwise stated. Shaded bands in cross-correlation plots represent bootstrap 95% confidence intervals estimated by resampling cells. Boxplot elements (median, interquartile range, whisker definition) are specified in the corresponding figure legends.

Statistical tests: For comparisons between two groups (e.g., Active vs Inactive), distributions were compared using the two-sided Mann-Whitney U test. For radial intensity profiles, two-sided Mann-Whitney U tests were performed independently at each radius and *p* values were adjusted using the Benjamini-Hochberg procedure across radii (reported as *q* values). For seq-DNA/RNA/IF-FISH re-analysis, per-gene Δ*z* values were tested against zero using two-sided one-sample Wilcoxon signed-rank tests. For sci-mtChIL-seq re-analysis, deviations of log_2_(Active/Inactive) CPT ratios from zero were assessed using two-sided one-sample Wilcoxon signed-rank tests; *p* values were corrected for multiple testing using the Benjamini-Hochberg procedure across all target-by-gene-set comparisons. Associations between locus-centered SNAPtag signals and transcriptional readouts were quantified using Spearman rank correlation. For lead-lag analyses of time-lapse trajectories, significance of the lead-lag index was assessed using a within-cell circular time-shift permutation test; permutation *p* values were adjusted using Benjamini-Hochberg FDR correction across tested factors. For HDAC inhibitor time-lapse experiments, duty-cycle distributions were compared using two-sided Mann-Whitney U tests with Holm correction for multiple comparisons versus the DMSO control.

Significance thresholds and the mapping from asterisks to *p*/*q* values are specified in each figure legend.

