## Supplemental figures for "Minute-scale coupling of chromatin marks and transcriptional bursts"

**Title:**

**Supplementary information**

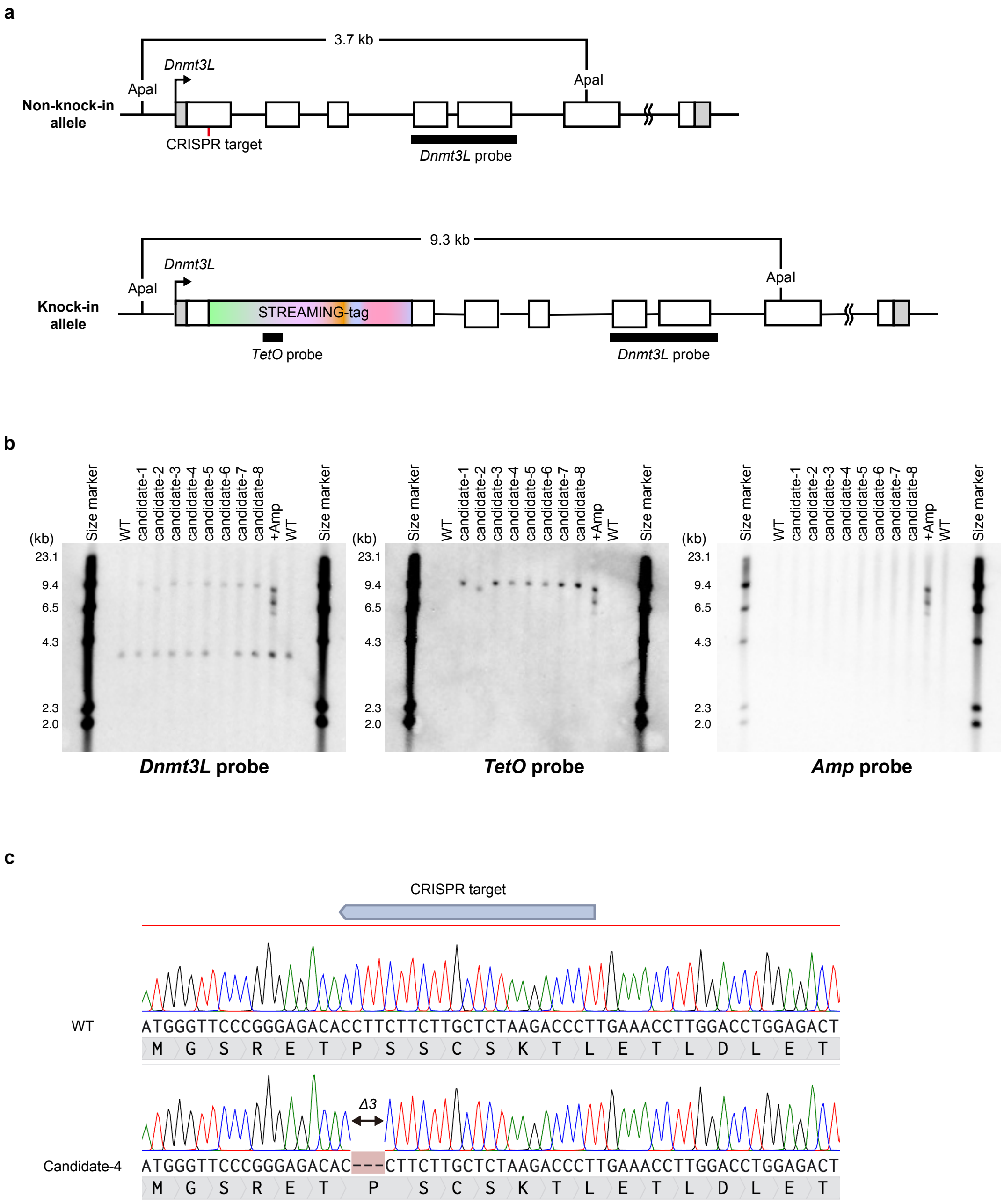


**Supplementary Fig. 1. Establishment of *Dnmt3L* STREAMING-tag knock-in cell lines. (a)** Gene structure of mouse *Dnmt3L* before and after STREAMING-tag knock-in. **(b)** Southern blot analysis of *Dnmt3L* STREAMING-tag knock-in candidate cells. **(c)** Sequence confirmation of non-knock-in allele in monoallelic knock-in candidate-4 cells. In this study, we used candidate-4 as a *Dnmt3L* STREAMING-tag knock-in cell.

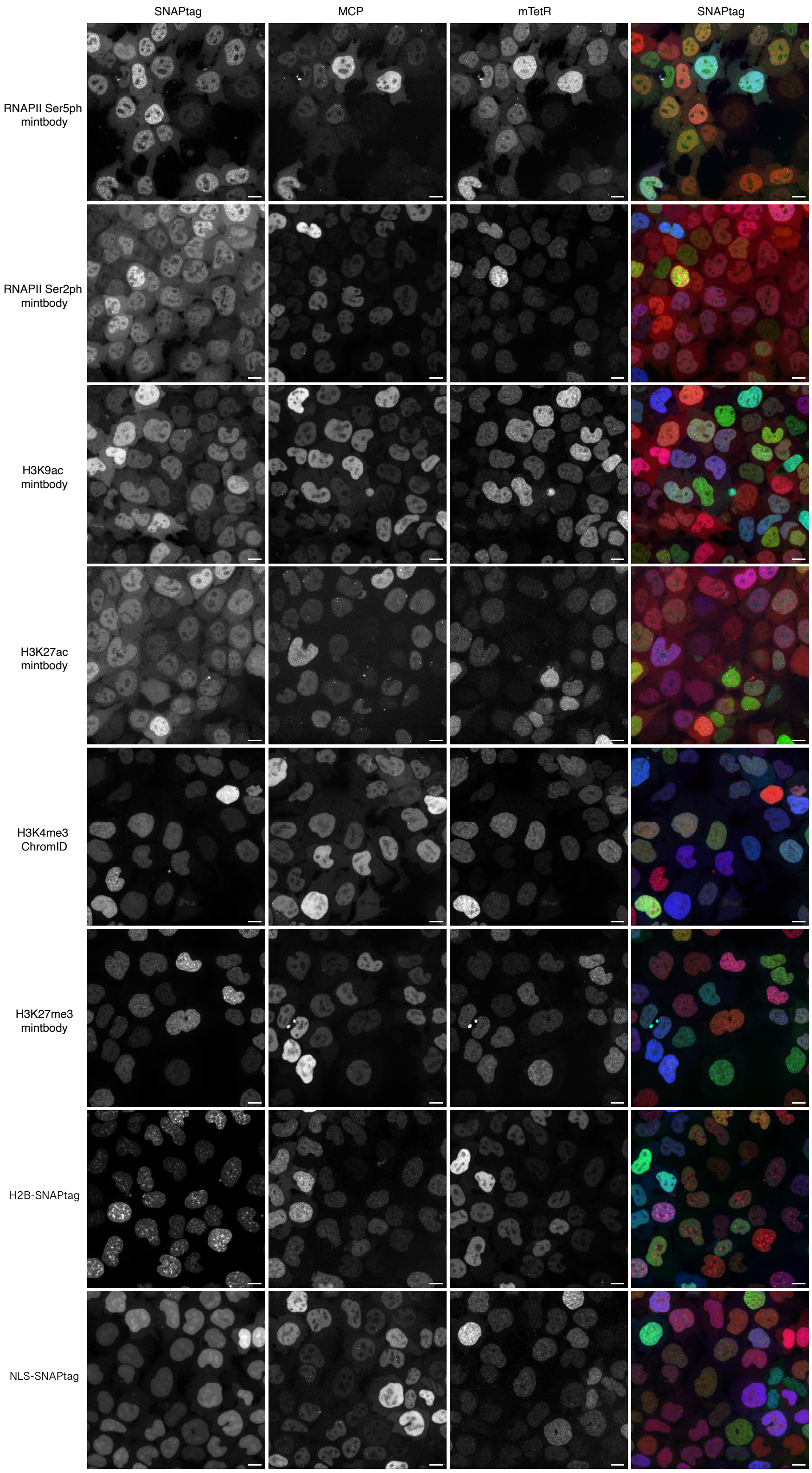


**Supplementary Fig. 2. Nuclear localizations of mintbody/ChromID probes.**

Fluorescence images of *Nanog* STREAMING-tag knockin cells expressing moderate levels of MCP-CFP and mTetR-YFP, together with SNAPtag-fused mintbody, ChromID, histone H2B, or a nuclear localization signal (NLS). Cells were imaged by confocal microscopy, and maximum intensity projection images are shown. Scale bar, 10 μm.

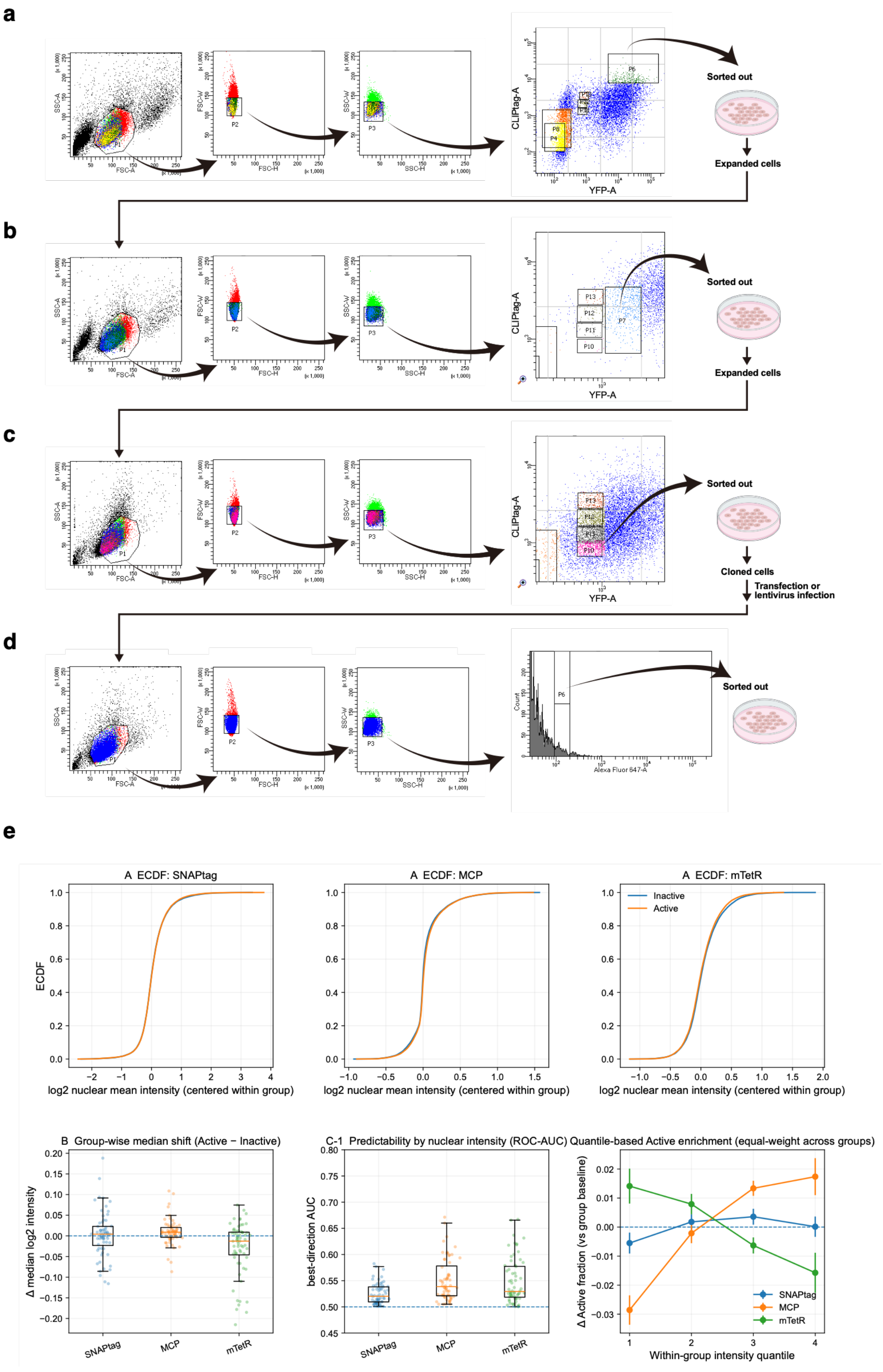


**Supplementary Fig. 3. Reporter nuclear mean intensities are weak predictors of transcriptional state calls. (a–c)** Fluorescence-activated cell sorting (FACS) strategy used to enrich and establish stable expression of the mTetR-YFP and MCP-CFP reporters. Because CFP fluorescence could not be detected by our flow cytometer, MCP expression was inferred using an NLS-CLIPtag co-expressed from the same transcript via a 2A peptide and labelled with CLIP-Cell TMR-Star. **(a)** First sort after transfection: high YFP and CLIPtag (P6). **(b)** Second sort after expansion: moderate YFP and CLIPtag (P7). **(c)** Third sort after further expansion: moderate YFP and CLIPtag (P10). Multiple rounds were performed because cells expressing only YFP or only CLIPtag can be abundant within the broad intermediate gate; repeated sorting enriches for double-positive cells with balanced expression. **(d)** After establishment of the reporter line, cells were transfected or lentivirally transduced to express SNAPtag fusion proteins and were sorted to obtain an appropriate SNAPtag expression range for imaging (example gate shown; P6). **(e)** For each nucleus/locus instance, nuclear mean intensities of SNAPtag, MCP and mTetR were extracted, log2-transformed, and centred within each imaging group (gene × factor/probe × replicate × well) by subtracting the group median to remove between-group offsets. Empirical cumulative distribution functions (ECDFs) of centred nuclear mean intensities for loci classified as Active or Inactive states were computed (pooled with equal weight across groups by subsampling an equal number of observations per group and state). Group-wise median shift between Active and Inactive states (Δ median log2 intensity), predictability of transcriptional state from nuclear mean intensity within each group quantified as best-direction ROC-AUC (max[AUC, 1−AUC]; 0.5 indicates chance-level prediction), and quantile-based enrichment of Active calls are shown as indicated.

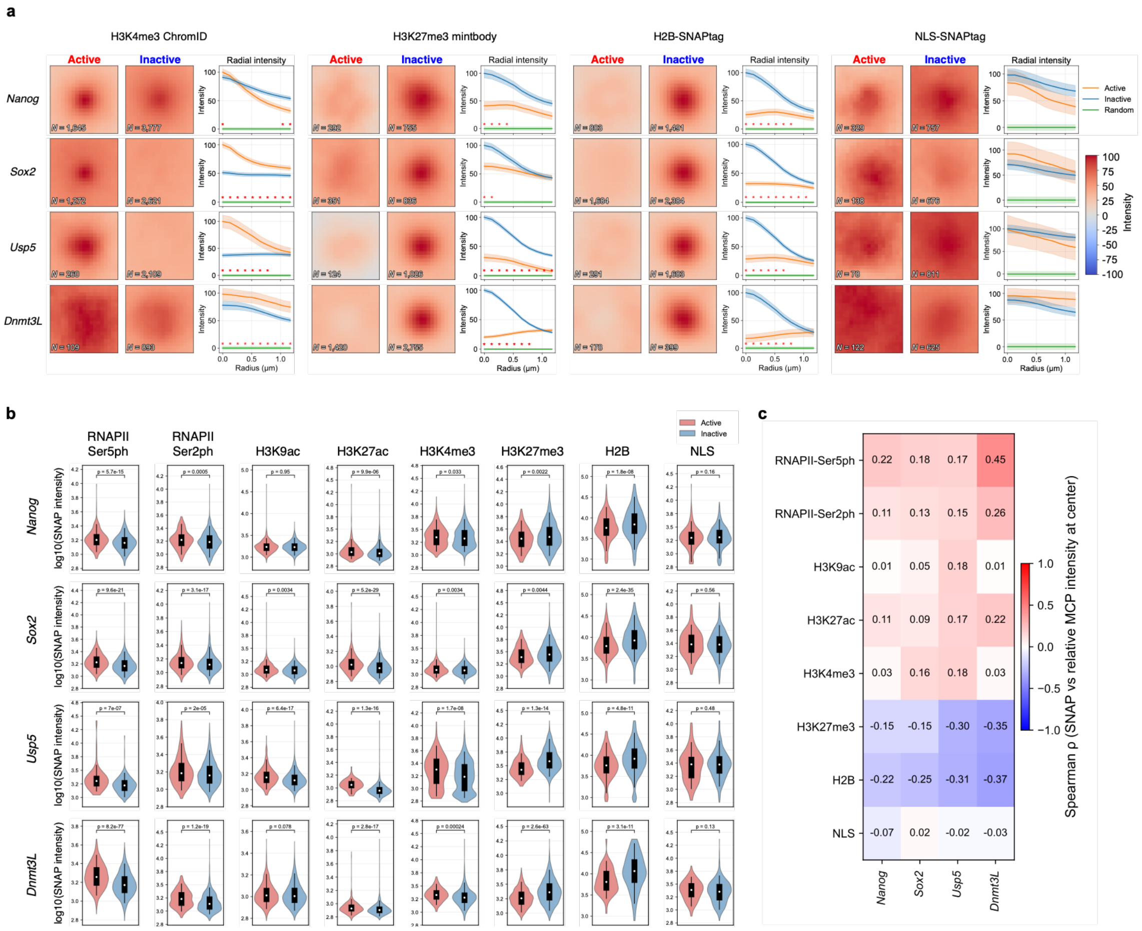


**Supplementary Fig. 4. Locus-centered histone-modification signals and their variability around genes during the transcriptional-burst cycle. (a)** Quantitative analysis of locus-centered signals reported by SNAPtag probes, including histone mark probes and the reference reporters H2B and NLS. For each locus instance, a 19 × 19-pixel crop (0.13 μm per pixel) centered on the mTetR focus (z-plane of the focus) was extracted from each channel and grouped by transcriptional state (Active versus Inactive). Pixel-wise mean locus-centered images were computed for each state after subtracting the mean image of size-matched random nuclear control crops sampled away from the locus. Radial intensity profiles show the mean ± s.e.m. across locus instances of the background-subtracted intensity in concentric 1-pixel annuli around the center. Red asterisks indicate radii with significant differences between Active and Inactive states (two-sided Mann-Whitney U test at each radius with Benjamini-Hochberg correction across radii; $q < 0.05$). **(b)** Distribution of SNAPtag fluorescence intensities at the center pixel of images centered on the gene locus. The distributions of fluorescence intensities of mintbody, ChromID, H2B, or NLS-SNAPtag at the pixel centered on the target gene (indicated on the left side of each panel) are shown. *P* values determined by the two-sided Mann-Whitney U test are indicated. In the boxplots, white dots represent the median, the upper and lower edges of the box represent the 25th and 75th percentiles, respectively, and the upper and lower ends of the black whiskers indicate the minimum and maximum values within the range from Q1 - 1.5×IQR to Q3 + 1.5×IQR. **(c)** Distribution of SNAPtag fluorescence intensities at the center pixel of images centered on the gene locus and relative MCP fluorescence intensities. For pixels centered on the target gene (indicated on the left side of each panel), the fluorescence intensities of mintbody, ChromID, H2B, or NLS-SNAPtag were calculated, along with the ratio between the mean fluorescence intensity within a region of radius 3 pixels from the center pixel and the mean fluorescence intensity of the entire 19 × 19 pixel image centered on the gene. Spearman correlation coefficients between these values were then calculated.

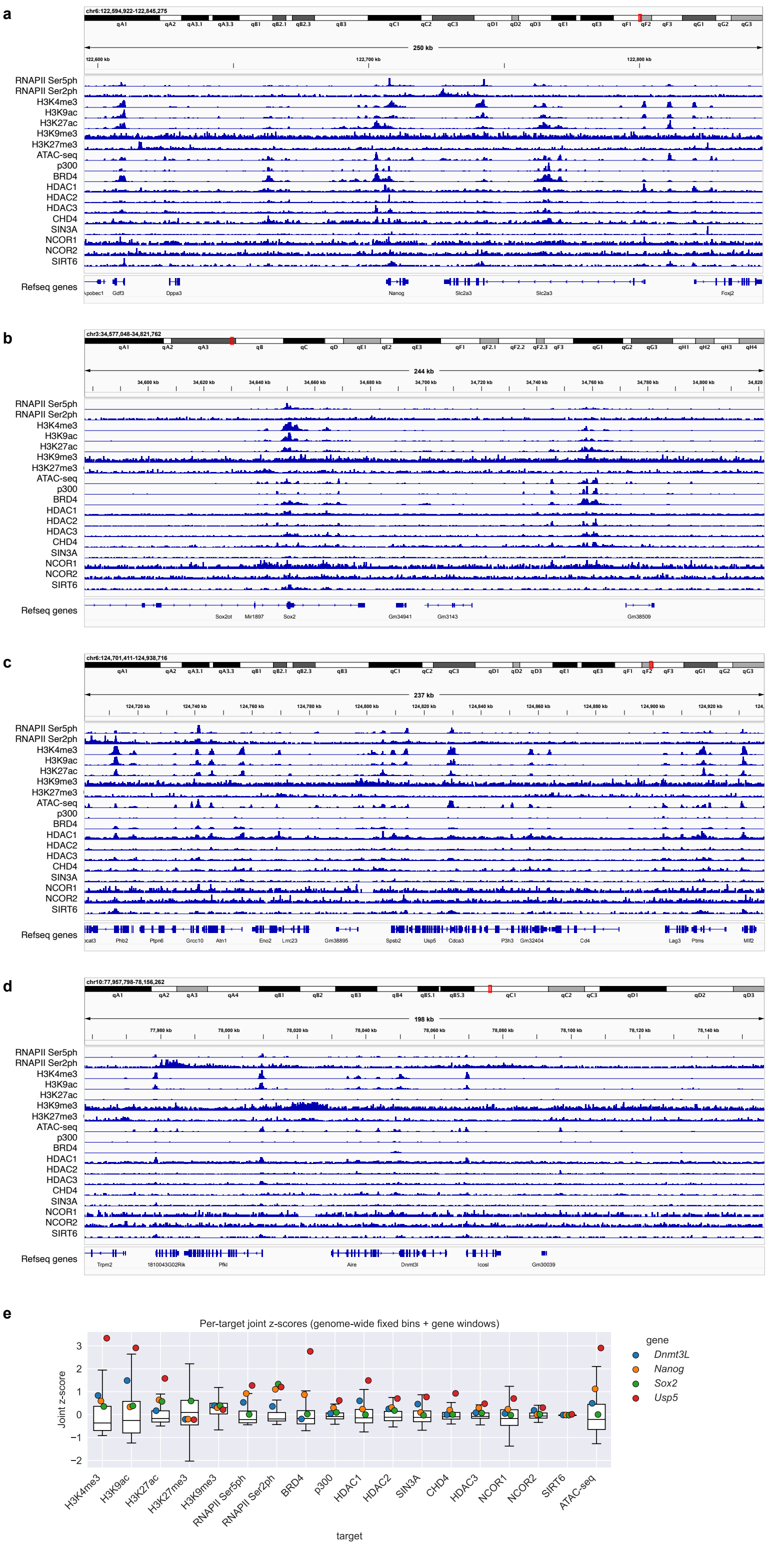


**Supplementary Fig. 5. Genome-wide context of chromatin marks and transcription-associated factors at loci analyzed by STREAMING-tag. (a-d)** Genome browser views (mm10) around the *Nanog* (a), *Sox2* (b), *Usp5* (c), and *Dnmt3L*

(d) loci showing representative ChIP-seq/ATAC-seq signal tracks for RNAPII CTD Ser5 phosphorylation (RNAPII Ser5ph) and Ser2 phosphorylation (RNAPII Ser2ph), histone modifications (H3K4me3, H3K9ac, H3K27ac, H3K9me3, H3K27me3), chromatin accessibility (ATAC-seq), and selected transcription-associated factors/cofactors and HDAC complex components (EP300/p300, BRD4, HDAC1/2/3, CHD4, SIN3A, NCOR1/2, and SIRT6). **(e)** Per-target joint z-scores comparing gene-centered windows to the genome-wide background distribution. For each target track, the mm10 genome was partitioned into non-overlapping fixed bins (0.5 Mb), and the mean bigWig signal was computed for each bin. For each gene, a 0.5-Mb TSS-centered window was defined and its mean signal was extracted. The 0.5-Mb scale was chosen to match the genomic segmentation used in related seq-DNA/RNA/IF-FISH analyses, and because locus-associated IF readouts have been shown to correlate with ChIP-seq enrichment when summarized at megabase-scale resolution[25]. For each target, signals were standardized using the combined distribution of genome-wide bin values and the gene-window values, yielding a joint z-score $z_{joint} = (x - \mu_t)/\sigma_t$, where $\mu_t$ and $\sigma_t$ are the mean and standard deviation of the combined values for target $t$. Boxplots show the distribution of $z_{joint}$ across genome-wide bins for each target, and colored points indicate $z_{joint}$ for each gene window (*Nanog*, *Sox2*, *Usp5*, and *Dnmt3L*). Accessions and file identifiers are listed in Supplementary Table 2.

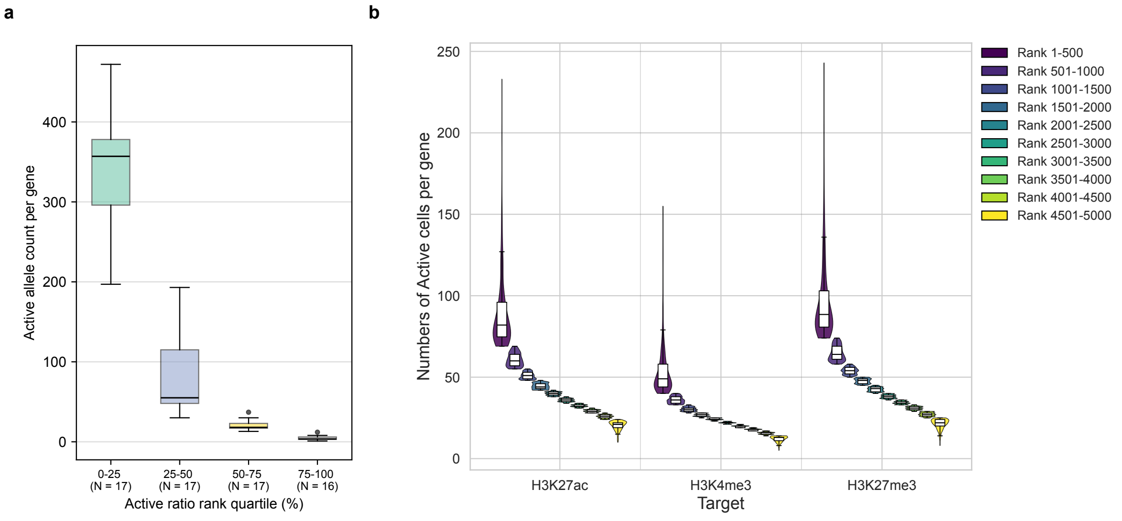


**Supplementary Fig. 6. Numbers of Active alleles or Active cells across gene-activity bins in orthogonal fixed-cell reanalyses. (a)** Numbers of Active alleles per gene in the seq-DNA/RNA/IF-FISH reanalysis (same gene ranking/stratification as in Fig. 2b). Box plots indicate median and interquartile range (IQR); whiskers denote 1.5× IQR. Points represent individual genes. **(b)** Numbers of Active cells per gene across activity-frequency bins in the sci-mtChIL-seq reanalysis. For each RNAPII Ser2ph–paired target dataset, the number of cells classified as Active state for each gene was computed. Violin plots with embedded box plots summarize the distribution across genes within each 500-gene bin.

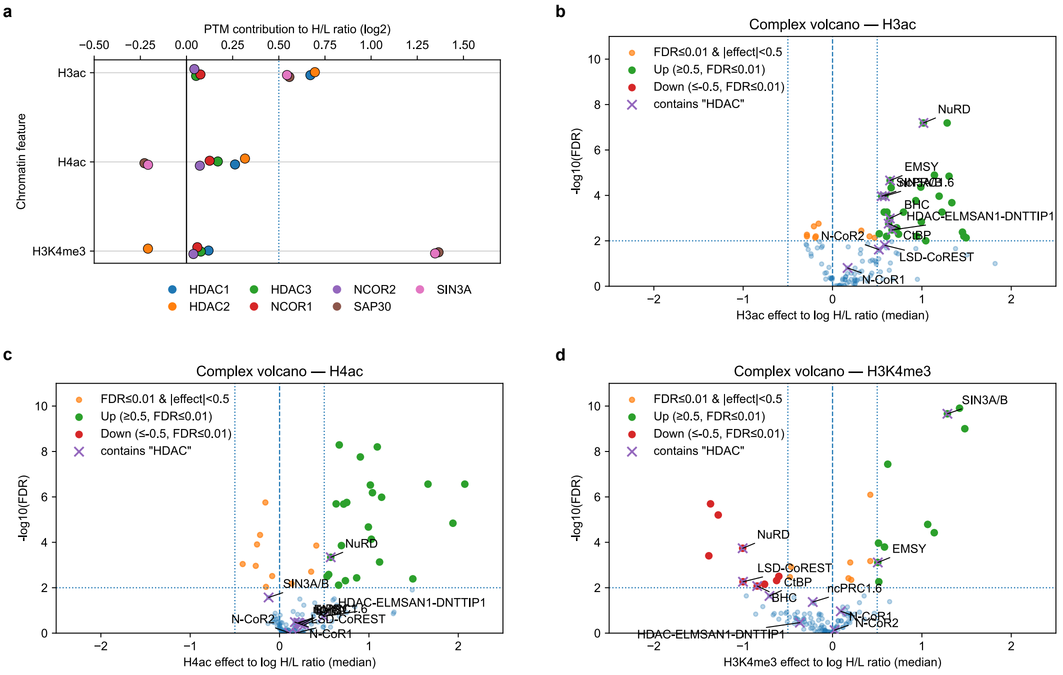


**Supplementary Fig. 7. Chromatin-binding properties of HDAC-related factors from re-analysis of MARCS data. (a)** Binding preferences of components of the NCOR2 complex (corum:1505; https://mips.helmholtz-muenchen.de/corum/?complex_id=1505), including HDAC1/2/3, NCOR1, NCOR2, SAP30, and SIN3A to H3ac, H4ac, and H3Kme3. **(b)** Volcano plot showing complex-wise binding to H3ac (complexes containing an HDAC are labeled). **(c)** Volcano plot for H4ac. **(d)** Volcano plot for H3K4me3 (complexes containing an HDAC are labeled).

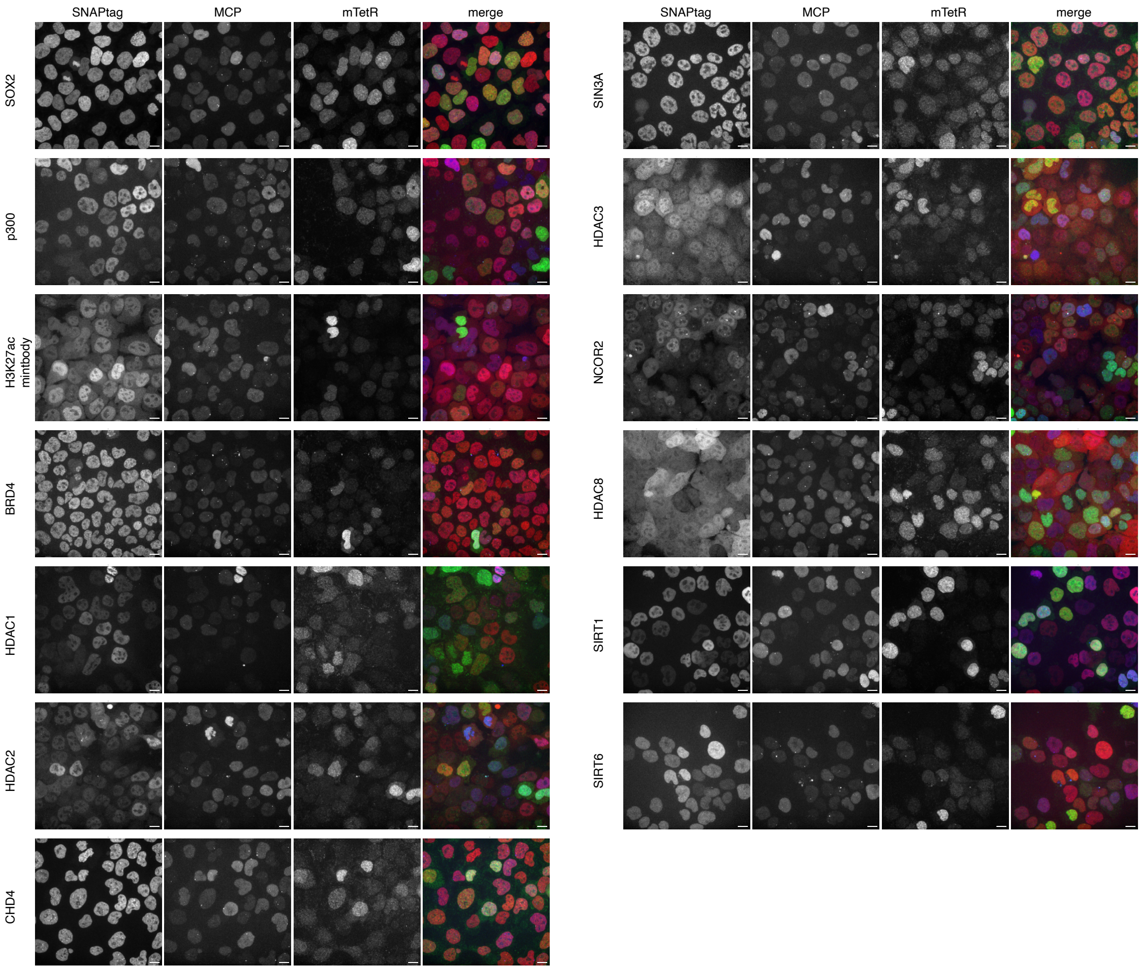


**Supplementary Fig. 8. Nuclear localizations of Mintbody/ChromID probes.**

Fluorescence images of *Nanog* STREAMING-tag knockin cells expressing moderate levels of MCP-RFP and mTetR-GFP, together with SNAPtag-fused proteins or H3K27ac mintbody. Cells were imaged by confocal microscopy, and maximum intensity projection images are shown. Scale bar, 10 μm.

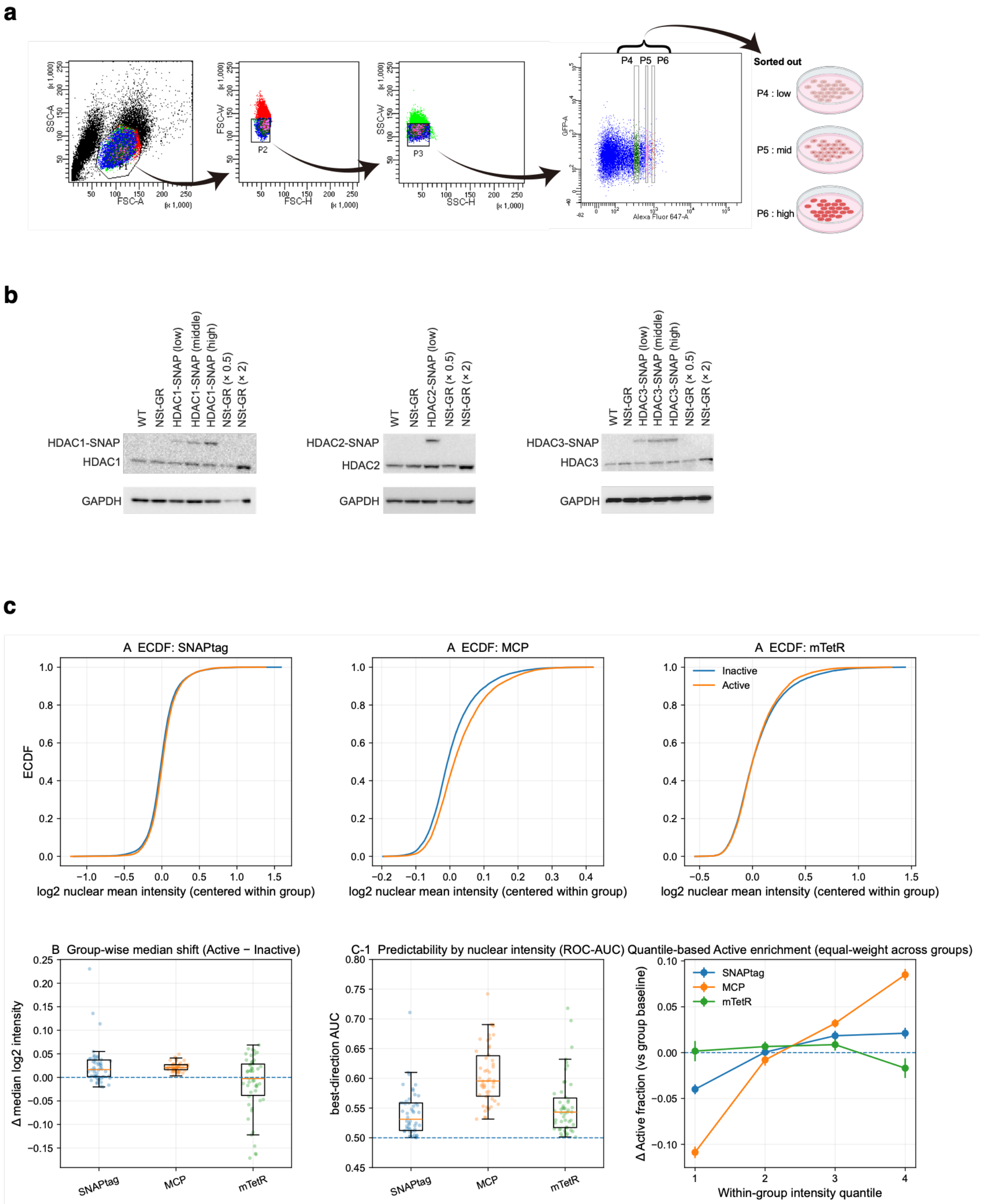


**Supplementary Fig. 9. Reporter nuclear mean intensities are weak predictors of transcriptional state calls. (a)** FACS gating strategy for SNAPtag fusion expression in the mTetR-GFP / MCP-RFP reporter background after transient transfection or lentiviral transduction. Cells were sorted into low (P4), mid (P5) and high (P6) SNAPtag-expression bins based on SNAPtag-ligand fluorescence. **(b)** Western blot analysis of HDAC1/2/3–SNAPtag expression in cells sorted from the gates in (a), compared with WT and parental NSt-GR cells (a cell line derived from *Nanog*

STREAMING-tag knockin cells moderately expressing MCP-RFP and TetR-GFP. See Supplementary Table 1). GAPDH, loading control. The low-expression bin (P4) yielded exogenous HDAC–SNAPtag levels comparable to or below endogenous HDACs and was therefore used for subsequent imaging experiments to minimize perturbation.

**(c)** Nuclear mean-intensity analyses of SNAPtag, MCP and mTetR signals (ECDFs, Active–Inactive median shifts, within-group ROC-AUC predictability, and quantile-based enrichment of Active calls), performed as in Supplementary Fig. 3e.
